# Satellite microglia gate neuronal excitability by shielding inhibitory synapses in adult brain

**DOI:** 10.64898/2025.12.13.694073

**Authors:** Shunyi Zhao, Mengdi Fei, Jiaying Zheng, Trace A. Christensen, Dimitrios Kleidonas, Yue Liang, Fangfang Qi, Koichiro Haruwaka, Lingxiao Wang, Gregory A. Worrell, Aivi T. Nguyen, Long-Jun Wu

## Abstract

Microglia, the primary immune cells of the central nervous system, are known for sculpting excitatory neural circuits via dynamic processes. However, their role in modulating inhibitory synaptic connections in adulthood remains largely unexplored. In this study, we identified an underappreciated microglia subpopulation, satellite microglia with physical soma-soma contact with neurons, that reshape inhibitory circuit connectivity and gate neuronal activity by shielding inhibitory synapses in the adult mouse brain. Using chronic in vivo two-photon imaging in adult mice, we observed in real time that satellite microglia associations displaced perisomatic inhibitory synapses whereas their dissociation permitted inhibitory synaptogenesis. Redistribution of satellite microglia reorganized inhibitory connectivity, altered local neural networks and changed behaviors. These findings establish satellite microglia as key architects of inhibitory circuitry and suggest their broader role in neural plasticity in health and disease.

## Introduction

Synaptic plasticity refers to the functional and structural modification of neural connections over time. This fundamental process underlies neural network dynamics and complex brain functions (*1*). Microglia, the primary immune cells in the central nervous system, actively sense neuronal activity and modulate synaptic functions in health and disease (*2, 3*). Emerging evidence highlights that microglia utilize their dynamic processes to participate in synaptic pruning, synaptic stripping, and synaptogenesis, underscoring a multifaceted microglial contribution to synaptic plasticity (*3, 4*).

Satellite microglia are defined by their unique soma-soma interaction with neurons (*5*). This microglia subpopulation constitutes approximately 40% of cortical microglia in the homeostatic brain (*6, 7*). They are distributed across various brain regions and are associated with multiple neuronal subtypes (*6*). Most prior studies have focused on satellite microglia in pathological conditions, where soma-soma interactions are more commonly observed (*8–11*). These pathological interactions have been proposed to phagocytose neurons and to strip perisomatic synapses (*9, 11, 12*). By contrast, the physiological functions of satellite microglia remain largely unknown.

In this study, we combined chronic two-photon imaging with correlative electron microscopy to investigate how satellite microglia shape the plasticity of axosomatic synapses. We demonstrate that satellite microglia selectively remodel inhibitory connections, thereby increasing neuronal activity and altering behaviors. Our findings uncover a unique role of satellite microglia in regulating circuit function and tuning brain activity.

## Results

### Satellite microglia form stable soma-soma interaction with neurons

Satellite microglia were identified by the proximity of their soma to neuronal cell bodies under light microscopy (*5, 6*). To examine the ultrastructure of this interaction, we performed correlative light-electron microscopy in *Cx3cr1^GFP/+^* mice received intracranial injection of AAV-CaMKII-tdTomato to label cortical excitatory neurons (Fig. 1A). Brain sections were marked using near-infrared branding (NIRB) for precise re-identification of neurons and associated satellite microglia under serial block face scanning electron microscopy (SBF-SEM) (Fig. S1). SBF-SEM confirmed direct membrane-membrane contact between satellite microglia and neuronal soma (Fig. 1A). We found that satellite microglia constituted 42.05 ± 2.52% of cortical microglia, 32.82 ± 4.20% of striatal microglia, 23.06 ± 1.54% of hippocampal microglia, and 19.65 ± 1.52 % of thalamic microglia (Fig. 1B and Fig. S2A). In the mouse cortex, satellite microglia primarily associated with excitatory neurons (Fig. 1C; 32.54 ± 2.85%), and rarely with PV^+^ inhibitory neurons (Fig. 1D and E; 1.95 ± 0.55%). In the human temporal lobe, satellite microglia made up 36.58 ± 3.99% of microglia (Fig. 1B and Fig. S2B) and showed membrane-membrane contact with both pyramidal (Fig. 1F) and interneuronal somata (Fig. 1G).

**Fig. 1.**
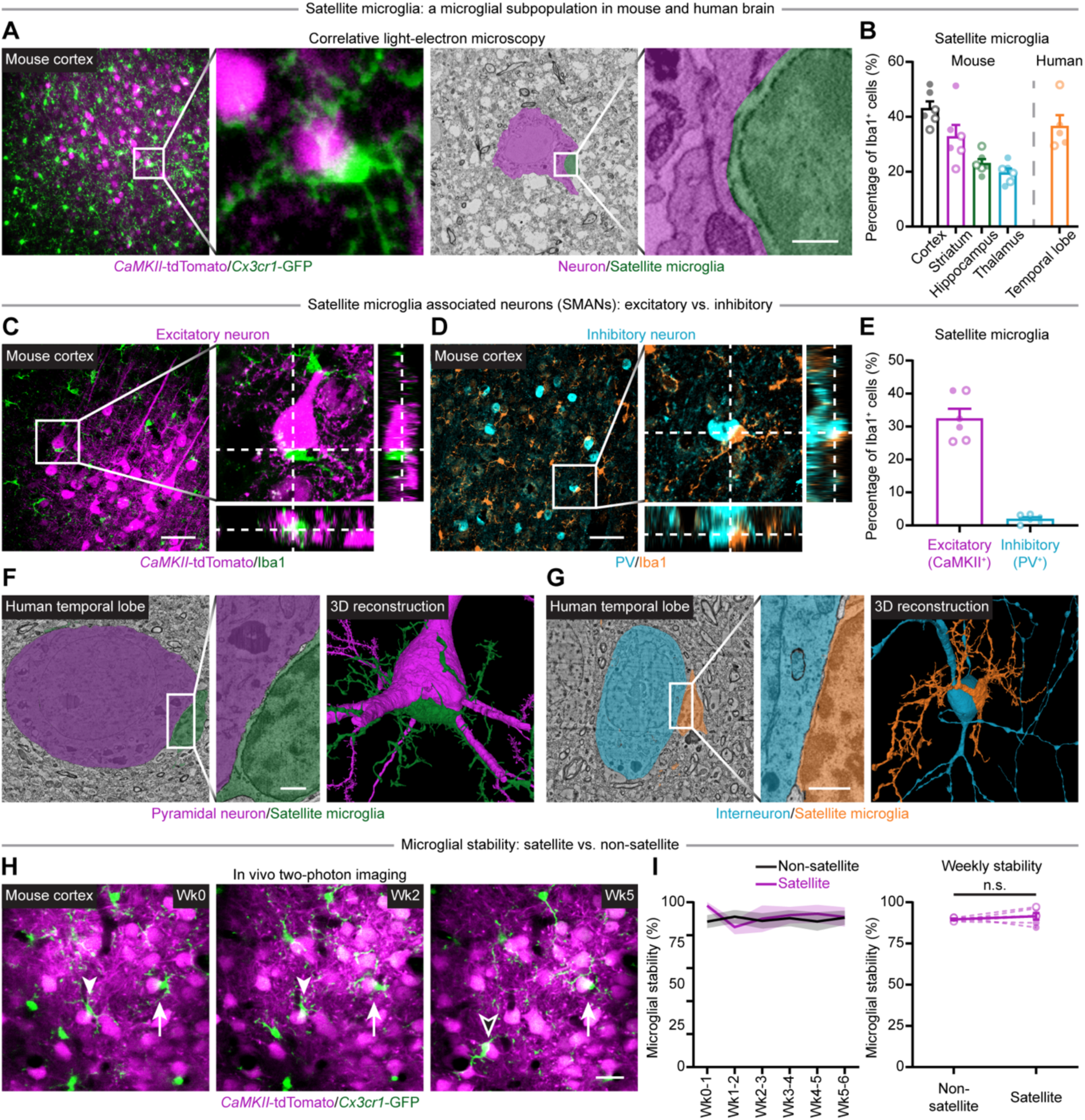
Satellite microglia-neuron soma interaction in mouse and human brains. (**A**) Two-photon imaging of satellite microglia (Cx3cr1^+^, green) and cortical excitatory neurons (CaMKII^+^, magenta) correlated with SBF-SEM images. (**B**) Quantification reveals that, satellite microglia constitute 42.05 ± 2.52% of cortical microglia. 32.82 ± 4.20% of striatal microglia, 23.06 ± 1.54% of hippocampal microglia, and 19.65 ± 1.52 % of thalamic microglia in the mouse brain (*N* = 6). In the human temporal lobe, satellite microglia constitute 36.58 ± 3.99% of microglia (*N* = 5). (**C** and **D**) Immunostaining illustrating satellite microglia interacting with excitatory neurons (C, CaMKII-tdTomato^+^, magenta) and inhibitory neurons (D, PV^+^, cyan) in the mouse cortex. (**E**) Quantification showing that 32.54 ± 2.85% of cortical microglia are satellite microglia associated with CaMKII^+^ excitatory neurons, and 1.95 ± 0.55% are associated with PV^+^ inhibitory neurons (*N* = 6). (**F** and **G**) Query of the H01 database illustrating the membrane-membrane contact between a satellite microglia and a pyramidal neuron soma (F) or an interneuronal soma (G) in the human temporal lobe. (**H** and **I**) In vivo two-photon imaging (H) demonstrating that satellite microglia (GFP, green) maintain stable soma-soma interaction (arrow) with neuronal cell bodies (tdTomato, magenta). Occasional dissociation (arrowhead) and formation (hollow arrowhead) of satellite microglia were observed. Quantification (I, *N* = 6) shows that satellite microglia have a high weekly stability rate (91.55 ± 2.17%), comparable with non-satellite microglia (89.60 ± 1.75%). Each point indicates an individual mouse or human subject (solid dots: male; hollow dots: female). Data are presented as mean ± SEM. Paired *t*-test (I). Scale bar: 1 μm (A, C, D), 20 μm (H), and 50 μm (E, F).

Homeostatic microglia have limited soma migration and dynamic process motility (*13, 14*). We next investigated the stability of satellite microglia using in vivo two-photon imaging (Fig. 1H). Tracking individual microglia over time, we found that satellite microglia consistently constituted approximately 40% of microglia (Fig. S3A). Additionally, their soma-soma interaction with neurons was remarkably stable, with a weekly stability rate of 91.55 ± 2.17% (Fig. 1I) and a turnover rate of 9.11 ± 2.33% (Fig. S3B), comparable with non-satellite microglia. Some satellite microglia remained associated with the same neuronal soma for over 18 weeks (Movie S1). In terms of process motility, satellite and non-satellite microglia showed similar process dynamics (Fig. S3C) and morphology (Fig. S3D), suggesting that both populations may have similar surveillance functions. Together, our data demonstrate that satellite microglia are a widespread subpopulation that engages in long-term soma-soma interaction with neurons.

### Satellite microglia are associated with higher neuronal activity

Recent studies have shown that microglia can regulate neuronal activity through their dynamic processes (*15–17*). To determine whether satellite microglia modulate neuronal activity via stable soma-soma interaction, we performed in vivo two-photon imaging to compare neuronal activity between satellite microglia associated neurons (SMANs) and non-SMANs. *Cx3cr1^GFP/+^* mice were intracranially injected with AAV-CaMKII-jRGECO1a for simultaneous imaging of microglia and calcium activity in cortical excitatory neurons (Fig. 2A). We found while both groups showed similar maximum calcium signal amplitude, SMANs had significantly higher active time and signal area compared to non-SMANs (Fig. 2B and Movie S2), indicating increased neuronal activity in SMANs.

**Fig. 2.**
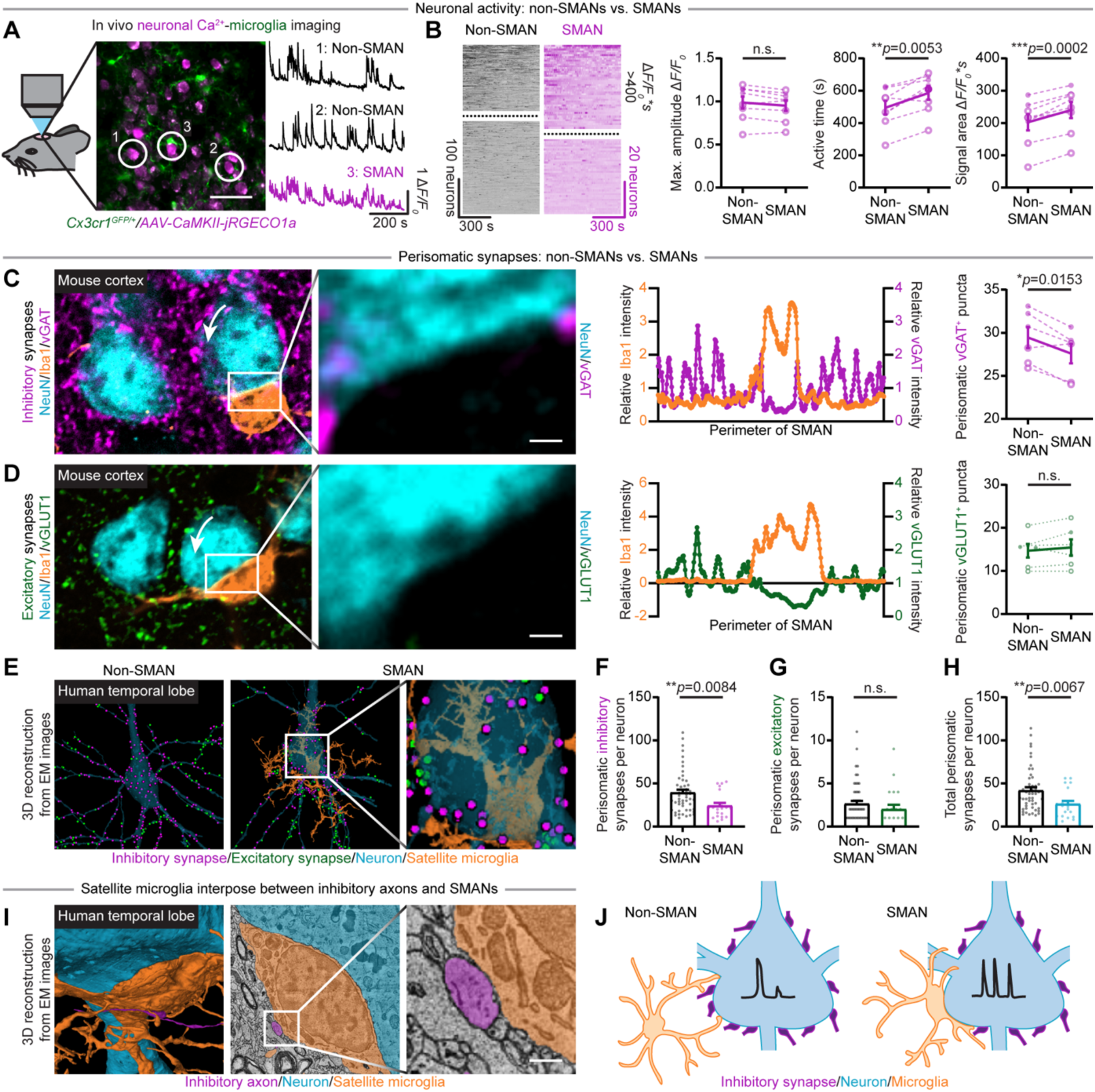
Satellite microglia-associated neurons (SMANs) show increased neuronal activity and reduced perisomatic inhibitory synapses. (**A**) Schematic of in vivo two-photon imaging strategy to identify satellite microglia (GFP, green) and evaluate neuronal activity (jRGECO1a, magenta). (**B**) Heatmaps and quantifications of Ca^2+^ activity for non-SMANs and SMANs. Dashed lines separate neurons with signal area above or below 400. Quantifications demonstrate similar maximum amplitude, but increased active time and signal area in SMANs (*N* = 8). Heatmap colors correspond to ΔF/F_0_ (−0.5 to 1.5), with darker shades indicate higher value. (**C**) Immunostaining and fluorescence intensity plot illustrating the absence of vGAT^+^ puncta (magenta) where satellite microglia (Iba1^+^, orange) contact neuronal soma (NeuN^+^, cyan). Quantification (*N* = 6) demonstrates reduced perisomatic inhibitory synapses around SMANs. (**D**) Immunostaining and fluorescence intensity plot illustrating the absence of vGLUT1^+^ puncta (green) where satellite microglia (Iba1^+^, orange) contact neuronal soma (NeuN^+^, cyan). However, quantification (*N* = 6) demonstrates comparable perisomatic excitatory synapses between SMAN and non-SMAN. (**E**-**H**) 3D reconstructions from the H01 EM dataset illustrating inhibitory (magenta) and excitatory (green) synapses on a non-SMAN and a SMAN (E). Inhibitory synapses are absent where satellite microglia (orange) contact neuronal soma. Quantifications reveal SMANs have fewer inhibitory (F) and similar excitatory (G) perisomatic synapses, leading to reduced total synapses (H). (**I**) 3D rendering and EM images from the H01 database showing satellite microglial soma (orange) interposed between neuronal soma (cyan) and inhibitory axons (magenta). (**J**) Diagram showing that SMANs have higher neuronal activity and reduced perisomatic inhibitory synapses. In (B-D), each point indicates an individual mouse (solid dots: male; hollow dots: female). In (F-H), each point indicates a single neuron. Data are presented as mean ± SEM. Paired *t*-test (B-D). Mann-Whitney test (F-H). n.s.: not significant. The exact *p* values are provided in the figure panels. Scale bar: 500 nm (I), 1 μm (C, D), and 50 μm (A).

Given that perisomatic synapses are predominantly inhibitory (*18*), we hypothesized that satellite microglia may occupy somatic space to limit perisomatic inhibition, thereby enhancing neuronal activity. Indeed, immunostaining of mouse cortex revealed an absence of synaptic markers at the contact site of satellite microglia (Fig. 2C and D). Quantification showed that SMANs were surrounded by fewer vGAT^+^ inhibitory synapses compared to non-SMANs (Fig. 2C). However, the number of excitatory synapses remained similar (Fig. 2D), likely because perisomatic excitatory synapses are relatively sparse. We also analyzed perisomatic inhibitory synapses surrounding SMANs and non-SMANs in human brain (H01 EM database) (Fig. 2E) (*19*). Human layer 5 pyramidal SMANs exhibited a reduced number of perisomatic inhibitory synapses (Fig. 2F) whereas the density of perisomatic excitatory synapses was unchanged (Fig. 2G). Consequently, the total number of perisomatic synaptic inputs was lower in SMANs (Fig. 2H). Altogether, both immunostaining and EM results suggest that satellite microglia reduce perisomatic inhibitory synapses on their associated neurons, which correlates with higher calcium activity in SMANs.

### Satellite microglia shield inhibitory synapses

While examining human brain EM images, we observed that satellite microglial somata also form direct membrane-membrane contacts with inhibitory axons (Fig. 2I). This close interaction among satellite microglia, neuronal somata and inhibitory axons suggests that satellite microglia may shield those perisomatic inhibitory synapses, regulating the structural plasticity of local circuits (Fig. 2J).

If satellite microglia shield inhibitory synapses, the dissociation of satellite microglia may open space for new inhibitory synapse formation (synaptogenesis), and their association could displace existing synapses (synaptic shielding). To test this hypothesis, we performed chronic in vivo two-photon imaging to directly observe the possible synaptogenesis or synaptic shielding. *PV^Cre/+^:Cx3cr1^GFP/+^* mice were injected with AAV-CaMKII-GCaMP6s and AAV-FLEX-tdTomato intracranially for identification of satellite microglia and imaging of axosomatic PV^+^ inputs (Fig. 3A). Satellite microglia were identified by the overlap between constant microglial GFP fluorescence and transient neuronal GCaMP6s signal (Fig. S4A), whereas non-SMANs showed only transient neuronal GCaMP6s signals (Fig. S4B). After observing synaptic remodeling, we performed correlative SBF-SEM to further examine the ultrastructure of the newly formed synapses (Fig. 3A).

**Fig. 3.**
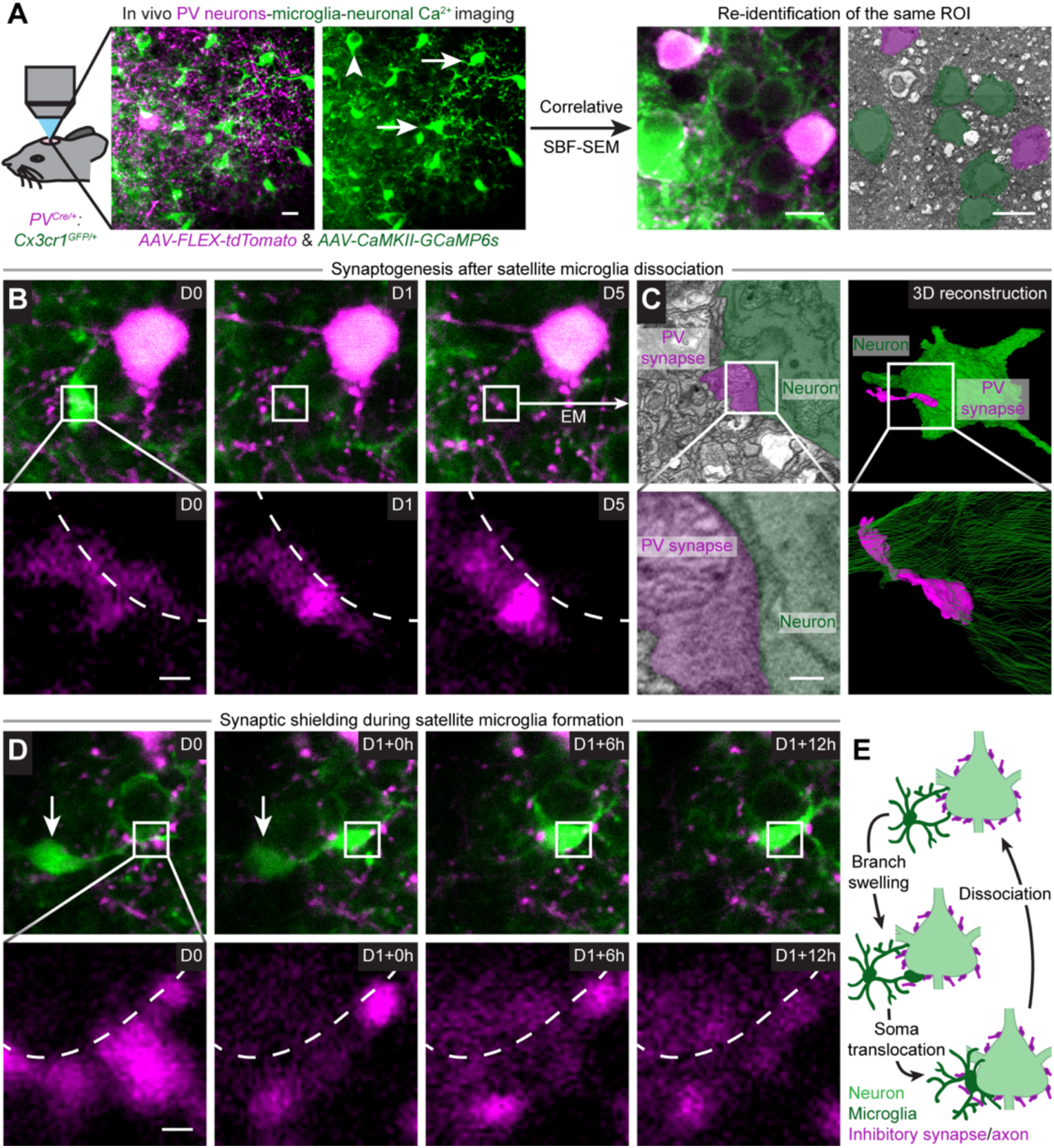
Satellite microglia participate in inhibitory synaptic plasticity by physically shielding neuronal somata. (**A**) Experimental design for correlative light-electron microscopy imaging. *PV^Cre/+^:Cx3cr1^GFP/+^* mice received AAV-FLEX-tdTomato and AAV-CaMKII-GCaMP6s injections to label PV^+^ neurons (tdTomato, magenta), excitatory neuronal somata (arrowhead; transient GCaMP6s signal, green), and microglia (arrow; constant GFP signal, green). ROIs identified by long-term two-photon imaging were marked by near-infrared branding to facilitate re-identification during SBF-SEM. High resolution EM images were acquired to examine the ultrastructure of newly formed PV^+^ puncta. (**B**) Time lapse two-photon imaging illustrating the formation of perisomatic PV^+^ synapses (magenta) following dissociation of satellite microglia (green). Satellite microglia were in contact with the neuronal soma on D0 but dissociated by D1. Dashed lines indicate neuronal somata. (**C**) SBF-SEM images and 3D reconstruction showing the ultrastructure of the newly formed perisomatic inhibitory synapse (corresponding to the boxed region in (B)). High resolution EM image reveals vesicle accumulation in the inhibitory synapse, suggesting active synaptic communication. (**D**) Time lapse two-photon imaging illustrating the displacement of perisomatic PV^+^ synapses (magenta) during satellite microglia (green) formation. The process begins with swelling of a microglial branch contacting the neuronal soma, followed by microglial soma translocation (arrow). Dashed lines indicate neuronal somata. (**E**) Diagram showing that satellite microglia actively modulate inhibitory synaptic plasticity by physically shielding neuronal somata. Scale bar: 200 nm (C), 1 μm (B, D), 10 μm (A).

Indeed, following satellite microglia dissociation, we observed the appearance of stable PV^+^ puncta at the contact site that had been occupied by satellite microglia. The newly generated PV^+^ inhibitory synapses remained stable during the rest of imaging time for days (Fig. 3B and Movie S3). Correlative SBF-SEM revealed that these new PV^+^ structures formed membrane-membrane contact with the neuronal soma and were enriched with synaptic vesicles, indicating the formation of functional inhibitory synapses (Fig. 3C and Movie S4).

Interestingly, we observed the formation of satellite microglia under in vivo imaging. During the formation of soma-soma contact, satellite microglia displaced axosomatic PV^+^ synapses (Fig. 3D and Movie S5). This process began with the swelling of a microglial branch, which partially displaced existing perisomatic PV^+^ synapses (Fig. 3D). Within 6 hours, the microglial soma migrated to the swelling site and displaced additional PV^+^ synapses (Fig. 3D). In summary, our results demonstrate that satellite microglia regulate the structural plasticity of perisomatic inhibitory synapses, facilitating synaptogenesis upon dissociation and contributing to synapse elimination during association (Fig. 3E).

### Redistribution of satellite microglia rewires neural connections

Do satellite microglia preferentially associate with a specific population of neurons? To address this question, we performed microglia ablation and repopulation to investigate whether repopulated microglia would reestablish soma-soma contact with the same neurons or adopt new neuronal partners. Mice were fed with PLX3397, a CSF1R inhibitor, for 2 weeks to efficiently ablate microglia. Afterward, normal chow was provided to allow microglia repopulation (Fig. 4A and B). We found that 7 days after switching to normal chow, amoeboid-shaped microglia began to appear and formed unstable soma-soma interaction with neurons (Fig. 4A and B). By day 14, microglia showed a ramified morphology, and satellite microglia became stable (Fig. 4C and Movie S6). Interestingly, only 14.73 ± 4.27% of former SMANs reestablished contact with repopulated satellite microglia. Moreover, 85.27 ± 4.27% of newly formed SMANs had not been previously associated with satellite microglia (Fig. 4D). These results suggest that SMANs do not represent a fixed neuronal subpopulation, and that satellite microglia redistribution may broadly reorganize neural connectivity via shielding inhibitory synapses.

**Fig. 4.**
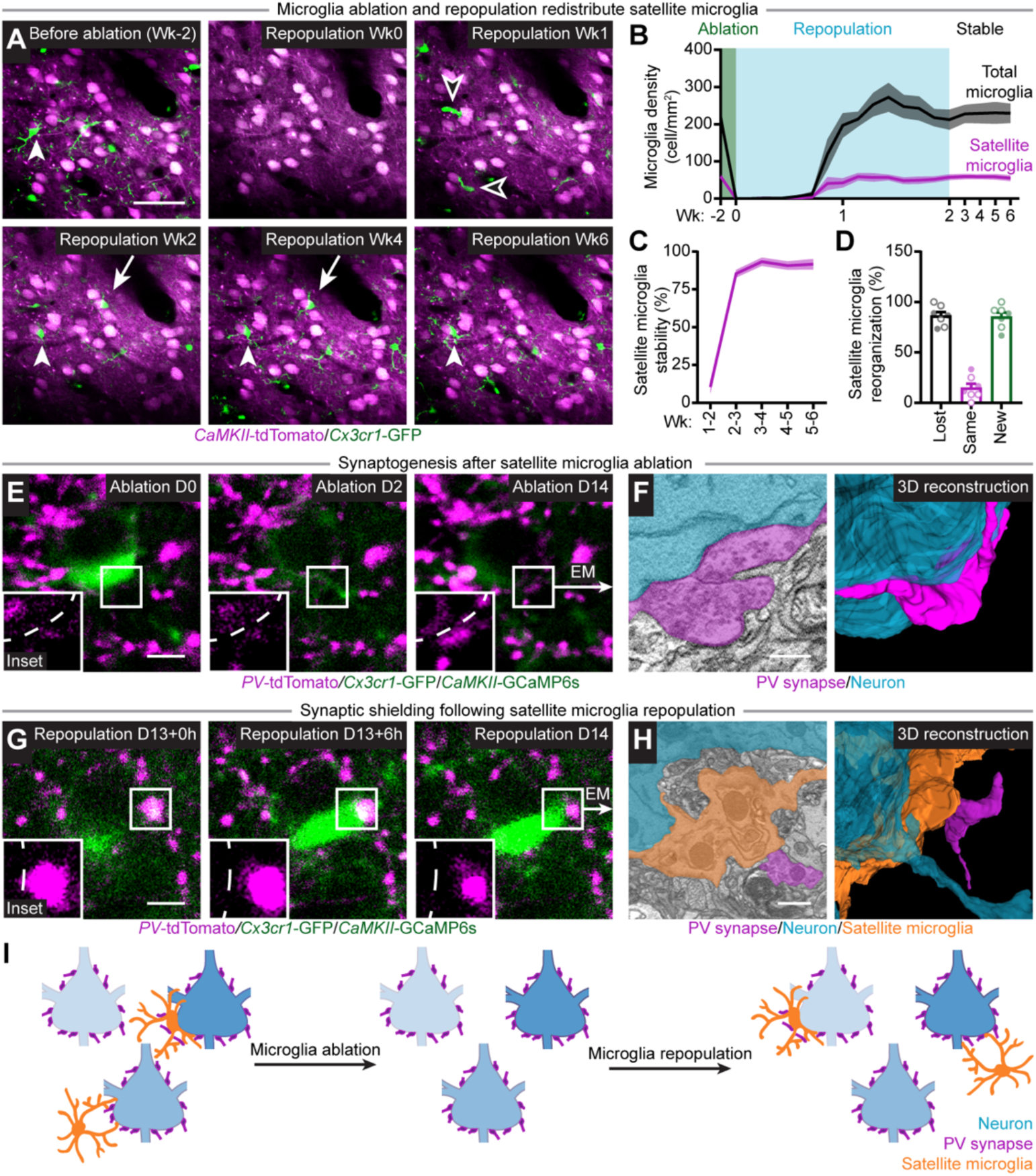
Satellite microglia redistribution rewires neural circuits. (**A**-**D**) Time lapse two-photon imaging illustrating the distribution of satellite microglia (green) before microglia ablation (Wk-2), after ablation (Wk0), during repopulation (Wk0-2), and following complete repopulation (Wk2-6) (A). Quantification (B) shows density changes of total and satellite microglia throughout the treatment. Satellite microglia were unstable during repopulation (A, hollow arrowhead) but became stable after complete repopulation (A, arrow and arrowhead) (C). Quantification (D) reveals that 86.22 ± 3.78% of SMANs lost satellite microglia following repopulation (lost), and 14.73 ± 4.27% obtained satellite microglia again (same; A, arrowhead). Additionally, 85.27 ± 4.27% of newly formed SMANs did not have satellite microglia before ablation (new; A, arrow). (**E**) Time lapse two-photon imaging illustrating the formation of perisomatic PV^+^ synapses (magenta) following microglia (green) ablation. Dashed lines indicate neuronal somata. (**F**) SBF-SEM image and 3D reconstruction showing the ultrastructure of the newly formed perisomatic inhibitory synapses (magenta; corresponding to the boxed region in (E)). Vesicle accumulation suggests active synaptic communication. (**G**) Time lapse two-photon imaging illustrating the displacement of perisomatic PV^+^ synapses (magenta) during microglia (green) repopulation (G). Dashed lines indicate neuronal somata. (**H**) SBF-SEM image and 3D reconstruction demonstrating the ultrastructure of newly formed satellite microglia (orange) and displaced inhibitory synapse (magenta, corresponding to the boxed region in (G)). High resolution EM image reveals satellite microglia (orange) interposed between the inhibitory axon (magenta) and the neuronal soma (cyan). (**I**) Diagram showing that experimental redistribution of satellite microglia modulates inhibitory synaptic plasticity by physically shielding neuronal somata. Scale bar: 500 nm (F), 1 μm (H), 5 μm (E, G), 50 μm (A).

To further examine neural rewiring, we performed in vivo two-photon imaging to visualize synapse formation and shielding during satellite microglia redistribution. Within several days of PLX chow treatment, we observed the loss of satellite microglia and the appearance of stable PV^+^ puncta at the previous satellite microglia contact site (Fig. 4E and Movie S7). Correlative SBF-SEM confirmed that the PV inputs had membrane-membrane contact with the neuronal soma and were enriched with synaptic vesicles (Fig. 4F and Movie S8), suggesting that the new inhibitory synapses were functionally active.

During early repopulation, we observed that unstable satellite microglia were still capable of displacing PV^+^ perisomatic synapses (Fig. S5A). Notably, when unstable satellite microglia migrated away, inhibitory synapses reestablished at the original position within hours (Movie S9), highlighting that synaptic shielding was rapidly reversible at this stage. Despite being transiently disconnected, reestablished inhibitory inputs retained synaptic vesicles (Fig. S5B and Movie S10), suggesting that active inhibitory synaptic transmission may be remained.

Following the complete microglia repopulation, redistributed stable satellite microglia displaced perisomatic inhibitory inputs by physically interposing between neuronal soma and inhibitory synapses (Fig. 4G and Movie S11). One day after displacement, the size of inhibitory synapse decreased (Fig. 4G), possibly suggesting ongoing synapse disassembly. Correlative SBF-SEM further confirmed that satellite microglia physically interposed between the neuronal soma and inhibitory synapses, forming membrane-membrane contacts with both structures (Fig. 4H and Movie S12). Overall, our chronic two-photon imaging and ultrastructural analysis indicate that satellite microglia redistribution can rewire local neural circuits by enabling inhibitory synaptogenesis upon ablation and shielding inhibitory synapses during repopulation (Fig. 4I).

### Redistribution of satellite microglia alters local neural network activity and changes behaviors

To investigate how redistribution of satellite microglia affects neural circuit function, we compared individual neuronal activity before and after microglia redistribution (Fig. 5A). Based on the association with satellite microglia, neurons were classified into 4 groups: (1) non-SMANs that remained as non-SMANs (non-SMANs), (2) SMANs that remained as SMANs (SMANs), (3) SMANs that lost their satellite microglia (former SMANs), and (4) non-SMANs that gained satellite microglia (newly formed SMANs).

**Fig. 5.**
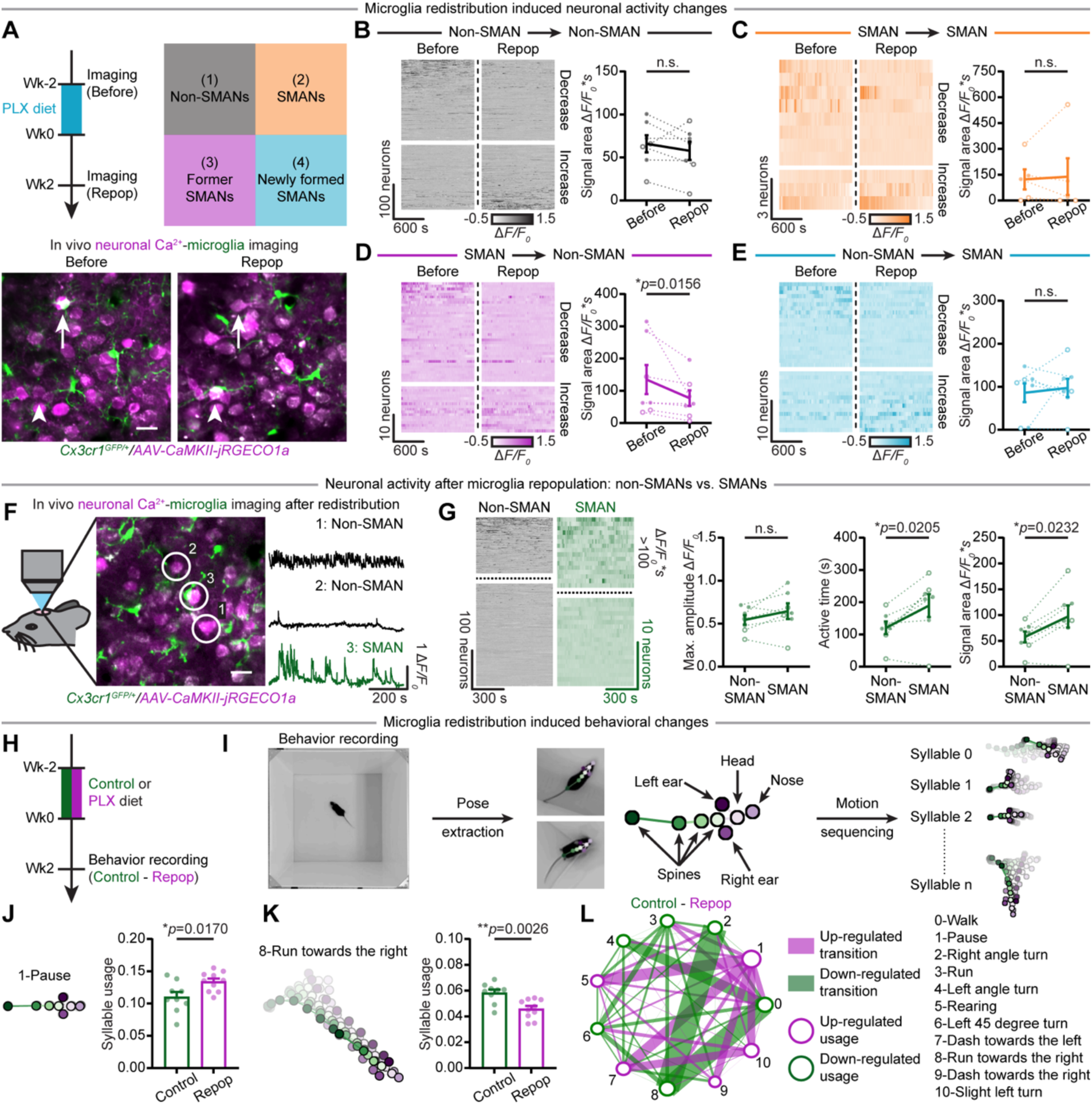
Redistribution of satellite microglia alters neuronal activity and behavioral patterns. (**A**) Timeline and representative two-photon images illustrating the strategy to evaluate neuronal activity after microglia redistribution. Based on the association with satellite microglia, neurons were classified into 4 groups: (1) non-SMANs (non-SMAN → non-SMAN; gray), (2) SMANs (SMAN → SMAN; orange), (3) former SMANs (SMAN → non-SMAN; magenta), and (4) newly formed SMANs (non-SMAN → SMAN; cyan). In two-photon images, arrows indicate former SMANs, and arrowheads indicate newly formed SMANs. (**B**-**E**) Heatmaps and quantifications of Ca^2+^ activity from non-SMANs (B), SMANs (C), former SMANs (D), and newly formed SMANs (E). Horizontal gaps separate neurons with increased or decreased activity. Dashed lines divide recordings taken before and after redistribution. Former SMANs showed decreased Ca^2+^ activity. (*N* = 7; *N* = 5 for SMANs, not detected in 2 mice) (**F**) Schematic of in vivo two-photon imaging strategy to identify satellite microglia (GFP, green) and compare neuronal activity (jRGECO1a, magenta) between SMANs and non-SMANs after microglia redistribution. (**G**) Heatmaps of Ca^2+^ activity for non-SMANs and newly formed SMANs following redistribution. Dashed lines separate neurons with signal area above or below 100. Quantifications show that newly formed SMANs have similar maximum amplitude, but increased active time and signal area (*N* = 7). Heatmap colors correspond to ΔF/F_0_ (−0.5 to 1.5), with darker shades indicate higher value. (**H** and **I**) Timeline (H) and schematic of Keypoint-MoSeq analysis (I). Mice were recorded in an open arena. Estimated poses were segmented into behavior motifs (syllables) (**J** and **K**) Pose trajectories (left) and quantifications (right) showing increased frequency of “pause” (J) and reduced frequency of “run towards the right” (K) following microglia redistribution. (**L**) Graph illustrating transition probability differences across syllables between groups. Edge thickness reflects the magnitude of difference. Each point indicates an individual mouse (solid dots: male; hollow dots: female). Data are presented as mean ± SEM. Paired *t*-test (B and G). Wilcoxon test (C-E). Multiple unpaired *t*-tests with FDR correction (*p* < 0.1; J and K). n.s.: not significant. The exact *p* values are provided in the figure panels. Scale bar: 20 μm (A and F).

We found that non-SMANs showed no significant changes in maximum amplitude, active time, and signal area (Fig. 5B and Fig. S6A). Similarly, SMANs that regained satellite microglia showed no significant change in neuronal calcium activity (Fig. 5C and Fig. S6B). In contrast, former SMANs showed decreased active time and signal area (Fig. 5D and Fig. S6C), suggesting reduced neuronal activity after losing satellite microglia (Movie S13). The newly formed SMANs had higher neuronal activity compared to non-SMANs in the repopulated brain, although they did not show statistically significant changes in calcium activity after microglia redistribution (Fig. 5E-G and Fig. S6D). These results suggest that satellite microglia may preferentially associate with hyperactive neurons during repopulation. Altogether, our findings suggest that satellite microglia redistribution leads to functional local circuit remodeling, especially for SMANs that lose satellite microglia.

To assess whether this neural circuit rewiring causes behavioral changes, mice were fed with control or PLX diet for 2 weeks, followed by 2 weeks of normal chow to allow microglia redistribution in PLX-treated mice (Fig. 5H). Then, we performed unsupervised behavioral analysis using keypoint tracking algorithms and motion sequencing (Keypoint-MoSeq) (*20*). Keypoint-MoSeq identifies mouse behavioral modules (syllables) and transition probabilities between syllables (Fig. 5I). Our results showed increased usage frequency of the “pause” syllable (Fig. 5J) and reduced usage frequency of the “run towards the right” syllable (Fig. 5K) in male mice with microglia redistribution. Other behaviors, such as “walk”, “right angle turn” and “Run”, remained comparable between the 2 groups (Fig. S7A). In addition to the differences in individual syllable usage, transitions between syllables were altered after complete repopulation, indicating disrupted behavioral flow (Fig. 5L). PLX-treated mice showed reduced transition probabilities among exploratory and locomotor syllables (“walk”, “right angle turn”, and “run towards the right”), and an increased probability of transitioning into “pause” (Fig. 5L). This pattern suggests that microglia redistribution changed movement flow. Mice were more likely to enter immobile states, instead of continuing exploration, after microglia redistribution. The behavioral alterations were more pronounced in male mice compared with female mice (Fig. S7B and C), potentially due to variability introduced by the estrous cycle (*21*).

## Discussion

Satellite microglia were first described by Dr. del Río-Hortega in 1919 (*5*). However, despite the surge of interest in microglial biology, satellite microglia have received less attention. Here, we demonstrate that satellite microglia actively govern the plasticity of perisomatic inhibitory synapses. Specifically, satellite microglia dissociation permits inhibitory synaptogenesis, while their association shields existing inhibitory synapses. Moreover, SMANs do not represent a fixed neuronal subpopulation. Instead, satellite microglia can be redistributed to different neurons, leading to inhibitory circuit rewiring, neuronal activity modulation, and behavioral changes. Altogether, our findings uncover a microglia-mediated synaptic plasticity, highlighting satellite microglia as a tractable entry point for regulating inhibitory circuits in health and disease.

Microglia remodel neural circuits through several mechanisms, including physical interactions with neurons (*15, 17, 22, 23*), synaptic pruning (*24–28*), extracellular matrix remodeling (*29*), and the release of soluble factors (*30, 31*). While most studies have focused on the regulation of excitatory synapses, recent work has begun to demonstrate microglial involvement in pruning inhibitory synapses during development (*25, 26*) and in diseases (*27, 28*). Our study reveals a distinct form of microglia-mediated inhibitory plasticity. The physical contact between satellite microglia and neurons limits the neuronal soma surface available for inhibitory synapse formation. When satellite microglia dissociate, the newly exposed surface permits inhibitory synaptogenesis. The exposed neuronal soma may engage nearby inhibitory axons via adhesion molecules and initiate intrinsic programs to recruit specific pre- and post-synaptic proteins for synaptogenesis (*32*). Alternatively, satellite microglia may release soluble factors upon dissociation to actively promote synaptogenesis.

In addition to facilitating inhibitory synaptogenesis, we demonstrate that satellite microglia also contribute to the displacement of perisomatic inhibitory synapses in the homeostatic brain. The concept of microglia-mediated synaptic displacement was proposed in a nerve axotomy model in 1968. The hypothesis was based on EM images showing no synaptic structure at the satellite microglia contact site (*33*). Since then, subsequent studies have indirectly tested this hypothesis in various neurological diseases, including traumatic brain injury (*8*), epilepsy (*9, 10*), and LPS-induced neuroinflammation (*11*). Using immunostaining and EM imaging, these studies consistently reported the absence of synapses at the contact site. However, it was unclear whether satellite microglia actively displace perisomatic inhibitory synapses or merely occupy regions where synapses had degenerated or failed to form. Here, we used live in vivo imaging with correlative EM to provide direct evidence for synaptic displacement by satellite microglia, resolving a long-standing question. Notably, the synaptic displacement occurs in the healthy brain, underscoring its relevance under physiological conditions.

PV^+^ inhibitory basket cells preferentially innervate neuronal somata to mediate fast and phasic inhibition (*34*). They play a critical role in regulating rhythmic population synchrony (*18*). Because specific oscillatory rhythms are associated with distinct cognitive functions, rewiring perisomatic inhibitory synapses can alter the excitation/inhibition balance, network states, and cognitive processing (*35, 36*). Indeed, our results showed that redistributing satellite microglia reshapes perisomatic inhibitory synapses on SMANs, thereby modulating local neural network activity and behaviors. Notably, even without experimental manipulation, spontaneous dissociation and re-engagement of satellite microglia similarly tune inhibitory connectivity, suggesting that satellite microglia actively contribute to neural network dynamics in the homeostatic brain.

Inhibitory circuits are essential for maintaining balance with excitatory circuits and play a critical role in diverse behaviors and cognition (*37*). Dysfunction of inhibitory circuits has been implicated in psychiatric disorders and cognitive impairments associated with neurodegenerative diseases (*38, 39*). In our study, satellite microglia redistribution altered behaviors during exploration, suggesting that the resulting rewired inhibitory circuits have functional consequences. However, our behavior analysis might underestimate the full impact of this remodeling. Since SMANs do not represent a fixed neuronal population (*6*), repopulated satellite microglia may associate stochastically with neuronal somata, introducing inter-individual variability in circuit changes. Therefore, identification of the molecular cues that govern satellite microglia dissociation and reassociation will enable a more precise manipulation of inhibitory circuitry and its behavioral outputs.

Microglia constantly monitor neuronal activity and regulate neuronal functions through their highly motile processes (*16, 40–43*). Whether satellite microglia can specifically sense neuronal activity and drive activity-dependent remodeling remains an open question. After satellite microglia redistribution, we found that newly formed SMANs had higher activity than non-SMANs but showed only limited changes relative to their baseline activity. This pattern may indicate that microglia preferentially associate with more active neurons during repopulation, making additional activity increases difficult to detect due to the ceiling effect. In seizure models, extracellular ATP promotes satellite microglia formation (*9*), whereas the proinflammatory cytokine, IL-1β, suppresses it and impairs motor learning by blocking neuronal ATP release (*44*). However, these signals are typically associated with neuronal injury rather than physiological modulation (*45, 46*). Identification of additional molecular cues will be critical for guiding satellite microglia to associate or dissociate with specific neuronal subtypes. Such targeted manipulation may offer therapeutic approaches for tuning neural circuits in psychiatric disorders and for mitigating cognitive dysfunctions in neurodegenerative diseases.

In summary, our study demonstrates that underappreciated satellite microglia play a critical role in inhibitory synaptic plasticity. Experimentally redistributing these cells rewires neural connections, modulates neuronal activity, and alters behaviors. Our results provide the mechanistic basis for future studies to explore satellite microglial roles in diverse behaviors. Moreover, dysregulation of satellite microglia may contribute to psychological disorders or cognitive impairments in neurodegenerative diseases.

## Acknowledgments

We thank Dr. Daniel R. Berger (Harvard University) for his suggestions on identifying microglia and oligodendrocyte progenitor cells in the H01 database. We also thank Drs. Doo-Sup Choi (Mayo Clinic), John D. Fryer (Translational Genomics Research Institute), Richard M. Weinshilboum (Mayo Clinic), and members of the Wu lab for their insightful discussions and valuable feedback. We thank Mayo Clinic Electron Microscopy Core facility for experimental and technical support. During the preparation of this manuscript, ChatGPT-4o and ChatGPT-5 were used only to improve readability.

## Author contributions

Conceptualization: SZ, LJW

Formal analysis: SZ, MF, JZ

Investigation: SZ, MF, JZ, TAC, DK, YL, FQ, KH, LW

Resources: ATN

Visualization: SZ

Funding acquisition: LJW

Supervision: LJW, GAW

Writing – original draft: SZ, LJW

Writing – review & editing: SZ, LJW

## Data and materials availability

All data are available in the main text or the supplementary materials.

## Supplementary Materials

### Materials and Methods

#### Animals

Wild-type (C57BL/6J, 000664), *PV^Cre/Cre^* (B6.129P2-*Pvalb^tm1(cre)Arbr^*/J, 017320) and *Cx3cr1^GFP/GFP^* (B6.129P2(Cg)-*Cx3cr1^tm1Litt^/*J, 005582) mouse lines were acquired from the Jackson Laboratory, and subsequently bred at either Mayo Clinic or University of Texas Health Science Center at Houston (UThealth Houston). Both male and female adult mice were included in all experiments. Mice were housed on a 12-hour light/dark cycle and in a climate-controlled environment with ad libitum access to food and water. All experimental procedures were approved by the Institutional Animal Care and Use Committee (IACUC) of Mayo Clinic or UThealth Houston.

#### Cranial window surgeries and virus injections

To prepare for cranial window surgeries, mice received 0.2 mg/mL ibuprofen in their drinking water for 3 days prior to the procedure. Anesthesia was induced with 3% isoflurane and maintained at 1.5% during surgery. Mice were kept on a heating pad throughout the surgery.

After removal of the scalp to expose the skull, a craniotomy was performed to create a circular 4-mm-diameter window above the somatosensory cortex (centered at AP: −2.0; ML: +1.5). Following the craniotomy, AAVs were injected into the somatosensory cortex using a glass pipette connected to a micropump (Drummond Scientific). During the virus injection, the exposed brain surface was kept moist with sterile phosphate-buffered saline (PBS). AAV2/5.CaMKII.tdTomato (Neurophotonics, 661-aav5) was used to label excitatory neurons. AAV9.CaMKII.GCaMP6s (Addgene, 107790) or AAV2/9.CaMKII.jRGECO1a (BrainVTA, PT3349) were used to image excitatory neuronal calcium activity. AAV9.CAG.FLEX.tdTomato (Addgene, 28306) was used to label PV^+^ inhibitory neurons in *PV^Cre/+^:Cx3cr1^GFP/+^* mice. A total of 300 nL of AAVs were injected to target layer II/III neurons (DV: −0.3, 200 nL, 40 nL/min; DV: −0.2, 100 nL, 20 nL/min) with a 5-minute period for virus diffusion following each injection.

After virus injection, a sterile 5-mm glass coverslip (Warner Instruments) was implanted and affixed with dental cement (Tetric EvoFlow). The remaining skull was covered with iBond Total Etch glue (Heraeus) and cured with an LED Curing Light. Additional dental cement (Tetric EvoFlow) was applied again around the glass coverslip and cured. A custom-made head plate was then secured to the window using dental cement. Following the surgery, mice recovered on a heating pad and received 0.2 mg/mL ibuprofen in their drinking water for an additional 3 days.

#### Two-photon imaging

Mice were allowed to recover for at least four weeks following cranial window surgery and underwent a 3-day habituation period prior to imaging. Each mouse was head-fixed under a two-photon microscope (Scientifica) equipped with an air-lifted platform (NeuroTar). Two-photon imaging was performed using a microscope equipped with a Mai-Tai DeepSee laser (Spectra Physics) or a Chameleon Ti:Sapphire laser (Coherent) tuned to 940 nm. A 16x water immersion objective lens (Nikon) was used, with digital magnification (zoom 2-6) applied according to experimental requirements. Emitted fluorescence signals from GFP and GCaMP6s were passed through a 525/50 filter (Chroma), and signals from jRGECO1a and tdTomato were passed through a 630/75 filter (Chroma). Laser power was maintained at 60 mW or below to minimize phototoxicity.

Imaging in the somatosensory cortex was conducted at 100-300 μm beneath the pial surface. For comparing process dynamics of satellite microglia and non-satellite microglia, Z-stacked images were acquired at 20 μm thickness with a 2 μm step size (1 stack per minute, 1024 × 1024 pixels, 3 fields of view). For long-term imaging of satellite stability or redistribution, the same Z-stack parameters were used (20 μm thickness, 2 μm step size, 1024 × 1024 pixels, 3 fields of view). For imaging the plasticity of perisomatic inhibitory synapses, Z-stacks were collected at 40 μm thickness with a 1 μm step size (1024 × 1024 pixels, 3 fields of view). For comparisons of calcium activity between SMANs and non-SMANs, neuronal activity was imaged at a frame rate of 1 Hz with a resolution of 512 × 512 pixels for 30 minutes. 5 fields of view were acquired per mouse.

#### Near-infrared branding (NIRB) and serial block-face scanning election microscopy (SBF-SEM)

NIRB was used to generate fiducial marks that facilitate the re-identification of regions of interest under SBF-SEM (*47*). To validate membrane-membrane contact between satellite microglia and neuronal somata, NIRB was performed on post-fixed brain slices. *Cx3cr1^GFP/+^* mice received intracranial *AAV-CaMKII-tdTomato* injection to label cortical excitatory neurons. 4 weeks after virus injection, mice were anesthetized and swiftly decapitated to prepare coronal slices (100 μm thickness) in oxygenated artificial cerebrospinal fluid (aCSF, 126 mM NaCl, 2.5 mM KCl, 1 mM NaH_2_PO_4_, 26 mM NaHCO_3_, 10.5 mM glucose, 1.3 mM MgSO_4_, 2 mM CaCl_2_; 300-305 mOsM, 7.35-7.4 pH). Slices were post-fixed overnight in 4% paraformaldehyde (PFA) in 0.1 M PBS. Fixed brain slices were mounted in PBS on glass slides, and satellite microglia were identified using confocal microscopy. Identified regions were subsequently imaged and marked under a two-photon microscope. For generating fiducial marks, the laser was focused using digital magnification (zoom 99) and tuned to 800 nm at maximum power. Several NIRB fiducial marks were generated surrounding the satellite microglia of interest. Slices were then transferred to SBF-SEM fixative (2% PFA, 2% glutaraldehyde, 0.15 M cacodylate buffer, 2 mM CaCl_2_; 7.4 pH) and stored for at least 24 hours before SBF-SEM imaging.

For examining the ultrastructure of newly formed or displaced PV^+^ puncta, NIRB was similarly performed on fixed brain tissue. After long-term two-photon imaging, mice were perfused with 4% PFA in 0.1 M PBS. The regions of interest were re-identified under a two-photon microscope, and NIRB fiducial marks were created around regions of interest using an 800 nm laser at maximum power and zoom 99. Samples were then trimmed and transferred to SBF-SEM fixative for at least 24 hours.

NIRB marked samples were processed for SBF-SEM by first rinsing in 0.1 M cacodylate buffer, followed by incubation in 2% osmium tetroxide and 1.5% potassium ferrocyanide in 0.1 M cacodylate. Samples were then washed with double distilled water (ddH_2_O), incubated at 50 °C in 1% thiocarbohydrazide, and again incubated in 2% osmium tetroxide in ddH2O. After rinsing in ddH2O, samples were incubated in 2% uranyl acetate overnight. The next day, samples were washed in ddH2O, incubated with Walton’s lead aspartate, dehydrated through a series of ethanol solutions, and embedded in EMbed 812 resin (Electron Microscopy Sciences). Embedded samples were first trimmed to remove excess resin and mounted onto aluminum stubs using Epo-Tek adhesive (Electron Microscopy Sciences). Samples were then placed into an SEM (Volumescope 2, Thermo Fisher Scientific), and NIRB marks were identified through initial sectioning and imaging. Once the NIRB marks were located, surrounding excess tissue was carefully trimmed using a diamond trimming tool (DiATOME, Electron Microscopy Sciences).

Imaging was performed under high vacuum/low water conditions with a starting energy of 1.8 keV and a beam current of 0.10 nA. For the validation of membrane-membrane contact between satellite microglia and neuronal somata, sections were cut at 100 nm thickness and imaged at 15 nm resolution (15 nm x 15 nm × 100 nm). For examining the ultrastructure of newly formed or displaced PV^+^ puncta, 4-6 tiled sections were cut at 50 nm thickness and imaged at 4 nm resolution (4 nm x 4 nm x 50 nm).

#### Correlated reconstruction of SBF-SEM images

SBF-SEM image processing was performed using ImageJ plugins, including 2D stitching for tiling, StackReg for alignment, Gaussian Blur for smoothing, and TrackEM2 for visualization. Reconstruct and PyReconstruct were used for 3D rendering (*48*). Neurons of interest were identified based on their spatial distribution in two-photon reference images and manually segmented at 50nm or 100nm intervals. Inhibitory presynaptic structures were estimated based on tdTomato labeled PV^+^ puncta observed in two-photon microscopy. Perisomatic inhibitory synapses were identified by the presence of synaptic vesicles and one or more symmetric electron-dense junctions with a neuronal soma. Microglia were identified by the presence of dense heterochromatin throughout the nucleus (*49*) and validated using two-photon reference images. Microglial processes were subsequently traced across the SBF-SEM image stacks.

#### Immunofluorescence analysis of mouse brain sections

To prepare mouse brain samples for immunostaining, isoflurane anesthetized mice were transcardially perfused with 40 mL PBS followed by 40 mL of cold 10% formaldehyde in PBS (LabChem). The whole brain was harvested, post-fixed in 10% formaldehyde overnight at 4 °C, and cryoprotected in 30% sucrose in PBS for at least 48 hours prior to cryosectioning. Coronal free-floating sections (30 µm) were obtained using a cryostat (Leica or Precisionary).

For immunostaining, sections were rinsed 3 times in Tris-buffered saline (TBS) and blocked with TBS buffer containing 5% donkey serum and 0.4% Triton X-100 at room temperature for 1 hour. Sections were then incubated overnight at 4 °C with combinations of the following primary antibodies: mouse anti-NeuN (1:1000, Abcam, Ab104224), rabbit anti-Iba1 (1:1000, Abcam, Ab178847), guinea pig anti-vGAT (1:500, Synaptic Systems, 131004), guinea pig anti-vGLUT1 (1:500, Synaptic Systems, 135304), and goat anti-PV (1:500, Swant, PVG213). Afterward, sections were washed 3 times in TBS and incubated for 2 hours at room temperature in blocking buffer with appropriate secondary antibodies: (1:500, Alexa-Fluor 555/594/647 donkey anti-mouse, Invitrogen, A31570/A21203/A31571; 1:500, Alexa-Fluor 488/555/594 donkey anti-rabbit, Invitrogen, A21206/A31572/A21207; 1:500, Alexa-Fluor 488 donkey anti-goat, Invitrogen, A11055; 1:500, Alexa-Fluor 488 donkey anti-guinea pig, Jackson ImmunoResearch, 706-545-148). Following 3 additional washes, sections were mounted with DAPI Fluoromount-G mounting medium (SouthernBiotech). Fluorescent images were acquired using a laser scanning confocal microscope (LSM 980, Zeiss; or AXR, Nikon). Z-stacked images (10-20 μm) were obtained by a 40x objective lens with digital magnification (zoom 1-3, 1024 × 1024 pixels) based on experimental requirements. For quantifying the satellite microglia population, images were acquired with a 1 μm step size. For quantifying perisomatic synapses, a step size of 0.274 μm was used. 3 fields of view were collected per mouse. Image brightness and contrast adjustments were made in ImageJ (National Institutes of Health).

#### Immunofluorescence analysis of human brain sections

Formalin-fixed paraffin-embedded (FFPE) human tissue sections were obtained from the Mayo Clinic Tissues Registry. All related experiments were approved by the Mayo Clinic Institutional Review Board (IRB; ID:21-012340). Temporal lobe sections were obtained from patients with mesial temporal lobe focal intractable epilepsy who elected to undergo neurosurgical resection. Included cases had no known history of focal cortical dysplasia, primary CNS tumor, or metastatic brain disease. Tissue from 5 patients was analyzed (3 male and 2 female, ranging in age from 20 to 35 years at the time of resection).

For immunostaining, tissue sections were first deparaffinized in xylene and rehydrated in a series of ethanol solutions, followed by antigen retrieval in citrate-based unmasking buffer (Vector Laboratories) using a steamer for 30 minutes. To reduce autofluorescence, sections were incubated with 0.1 % Sudan Black for 30 minutes and rinsed with 70% ethanol, 50% ethanol and PBS. Following these steps, sections underwent immunostaining as previously described. Sections were incubated overnight at 4 °C with mouse anti-NeuN (1:500, Abcam, Ab104224) and rabbit anti-Iba1 (1:1000, Abcam, Ab178847) primary antibodies. The next day, sections were incubated with secondary antibodies (1:500, Alexa-Fluor 594 donkey anti-mouse, Invitrogen, A21203; 1:500, Alexa-Fluor 488 donkey anti-rabbit, Invitrogen, A21206). Fluorescent images were acquired using a laser scanning confocal microscope (LSM 980, Zeiss). Z-stacked images (5 μm) were obtained using a 40x objective lens (1024 × 1024 pixels) with a 1 μm step size. 3 fields of view were collected per human subject. Image brightness and contrast were adjusted using ImageJ (National Institutes of Health).

#### Microglia ablation

Microglial ablation was achieved through the administration of chow containing the colony-stimulating factor 1 receptor (CSF1R) inhibitor, PLX3397 (600 mg/kg, Chemgood), provided ad libitum for 2 weeks to ensure complete depletion of microglia.

#### Behavioral tests

Mice were transferred to the test room at least 1 hour prior to the start of behavioral measurements. Behavioral tests were performed during the light phase. To assess spontaneous locomotor activity, mice were placed in a square open-field arena (40 x 40 x 30 cm) and allowed to explore freely for 15 minutes. Mouse behavior was recorded by an overhead camera. Recorded videos were analyzed using DeepLabCut and Keypoint-MoSeq. DeepLabCut was used to estimate mouse posture and track multiple body parts, including the head, nose, left ear, right ear, 4 points along the spine, and the tail (*50*). Keypoint-MoSeq is an unsupervised behavioral analysis tool that utilizes keypoint tracking algorithms and motion sequencing to identify mouse behavioral modules (syllables) and transition probabilities between them (*20*). For Keypoint-MoSeq analysis, mouse tail position was excluded to decrease false syllable detection. Only syllables with a usage frequency greater than 0.03 were included in the statistical analysis. Each syllable was manually named by investigators after examining the corresponding video clips and trajectory plots.

#### Query of H01 EM dataset

Ultrastructure identification of satellite microglia and SMANs in the human temporal lobe was performed by querying the H01 EM database (*19*). Microglia were first identified either from a set of 27 pre-identified microglia examples or from the “microglia/OPC” group. Satellite microglia and SMANs were then identified based on membrane-membrane contact between microglia and neurons.

To quantify somatic inhibitory and excitatory inputs surrounding SMANs and non-SMANs, the pre-annotated “incoming excitatory” and “incoming inhibitory” layers were used, and synapse numbers were manually counted.

#### Characterization of satellite microglia

To quantify the percentage of satellite microglia, overlaps between microglial somata and neuronal somata were manually examined at each Z-level from two-photon or confocal images. Imaris was used to validate manually identified cortical satellite microglia from confocal images using the following parameters. (NeuN: [Algorithm]: Enable Region Of Interest = false, Enable Region Growing = true, Enable Tracking = false, Enable Classify = false, Enable Shortest Distance = true; [Source Channel]: Source Channel Index = 1, Enable Smooth = true, Surface Grain Size = 1.00 µm, Enable Eliminate Background = true, Diameter Of Largest Sphere = 10.0 µm; [Threshold]: Active Threshold = true, Enable Automatic Threshold = false, Manual Threshold Value = 28 (adjusted between staining batches), Active Threshold B = false, Region Growing Estimated Diameter = 5.00 µm, Region Growing Morphological Split = false; [Filter Seed Points]: “Quality” above 9.00; [Filter Surfaces]: “Number of Voxels Img=1” above 650. Iba1: [Algorithm]: Enable Region Of Interest = false, Enable Region Growing = false, Enable Tracking = false, Enable Classify = false, Enable Shortest Distance = true; [Source Channel]: Source Channel Index = 3, Enable Smooth = true, Surface Grain Size = 0.900 µm, Enable Eliminate Background = true, Diameter Of Largest Sphere = 8.00 µm; [Threshold]: Active Threshold = true, Enable Automatic Threshold = false, Manual Threshold Value = 48 (adjusted between staining batches), Active Threshold B = false; [Filter Surfaces]: “Number of Voxels Img=1” above 500.) Both manual and automated methods produced similar percentages. Therefore, manual criteria were applied in cases where Imaris failed to accurately reconstruct neuronal somata.

For quantifications of satellite microglia stability and turnover rate, longitudinal two-photon images acquired over several weeks were analyzed at each Z-level. The total number of microglia, satellite microglia, newly formed satellite microglia, and dissociated satellite microglia were recorded. Stability was calculated as the number of satellite microglia present at both time points (D_x_ to D_y_) divided by the number at D_x_. Turnover rate was defined as the number of satellite microglia lost between D_x_ and D_y_ divided by the average number of satellite microglia at D_x_ and D_y_.

To assess satellite microglia reorganization following microglial ablation and repopulation, two-photon images acquired before ablation and after complete repopulation were compared. “Lost” was defined as the proportion of SMANs that lost their satellite microglia after repopulation, relative to the SMAN count before ablation. “New” was defined as the proportion of non-SMANs that gained satellite microglia after repopulation, relative to the SMAN count after repopulation. “Same” was defined as the proportion of SMANs that remained SMANs after repopulation, relative to the SMAN count before ablation.

For microglia morphology analysis from Iba1 immunostaining, Z-projected images were segmented using the Trainable Weka Segmentation plug-in in ImageJ. 10 satellite microglia and 10 non-satellite microglia per mouse were randomly selected for Sholl analysis.

For analysis of microglia process motility in two-photon images, primary branch movements were manually tracked using the Manual Tracking plug-in in ImageJ. To compare process movement speed between non-satellite microglia and satellite microglia, at least 5 extending and 5 retracting primary branches were measured per cell.

#### Quantification of perisomatic synapses via immunostaining

To compare perisomatic synapses between SMANs and non-SMANs from confocal images, a segmented line (1 μm wide) was drawn along the NeuN^+^ neuronal soma in ImageJ, with the midline of the segmented line positioned at the soma edge. For each neuron, 3 Z-panels spaced 0.4 μm apart were analyzed. In each Z-plane, a single vGAT or vGLUT1 channel image was analyzed using the “Plot Profile” tool to generate fluorescence intensity profiles along the segmented lines. Raw fluorescence intensity values were converted into ΔF/F_0_, where F_0_ was defined as the 50^th^ percentile value for each segmented line. Perisomatic synapses were defined as regions where ΔF/F_0_ exceeded 0.5 for a continuous distance of more than 0.25 μm along the line. The number of perisomatic synapses surrounding SMANs and non-SMANs was then quantified. For this analysis, 5-14 SMANs and 30 non-SMANs per mouse were randomly selected.

#### Neuronal calcium imaging analysis

Two-photon time-lapse images underwent registration using the ImageJ plugin, TurboReg, to correct any image shifts. For the analysis of neuronal calcium activity, an average intensity image of the entire video was generated to facilitate the selection of neuron somata and the identification of SMANs. Neuron somata were manually drawn using the oval selection tool. Then, mean fluorescent intensity values were obtained and subsequently converted into ΔF/F_0_. The baseline fluorescence was defined as the lowest 25^th^ percentile value across all frames. Neurons exhibiting ΔF/F_0_ > 0.25 were considered “active”. Parameters such as neuronal active time and signal area were calculated accordingly.

#### Statistics

Detailed statistical information, including sample size and statistical methods, is provided in the figure legends corresponding to each specific experiment. Generally, normality was first assessed. For data that followed the Gaussian distribution, an unpaired 2-tailed *t-*test was employed for comparing experiments involving 2 groups. In instances where 2 time points or 2 groups of cells from the same animal were compared, a paired 2-tailed *t-*test was used. For unpaired groups that did not follow the Gaussian distribution, the Mann-Whitney test was used. For paired groups that did not follow the Gaussian distribution, the Wilcoxon test was used. For Keypoint-MoSeq analysis, multiple unpaired *t*-tests with FDR correction (*p* < 0.1) were performed to limit false positive results. Results are presented as the mean ± standard error of the mean (SEM), and statistical significance was determined when *p*<0.05. Statistical analyses were performed using GraphPad Prism 10 software. Experimental designs and sample sizes were determined to minimize animal usage and distress while ensuring sufficiency for detecting robust effect sizes. Researchers were aware of genotypes and treatments during experiments and data analysis. To mitigate potential bias, we implemented rigorous controls and automated data analysis wherever possible. These analytical steps are detailed in the Methods section to ensure transparency and reproducibility of our findings.

**Fig. S1.**
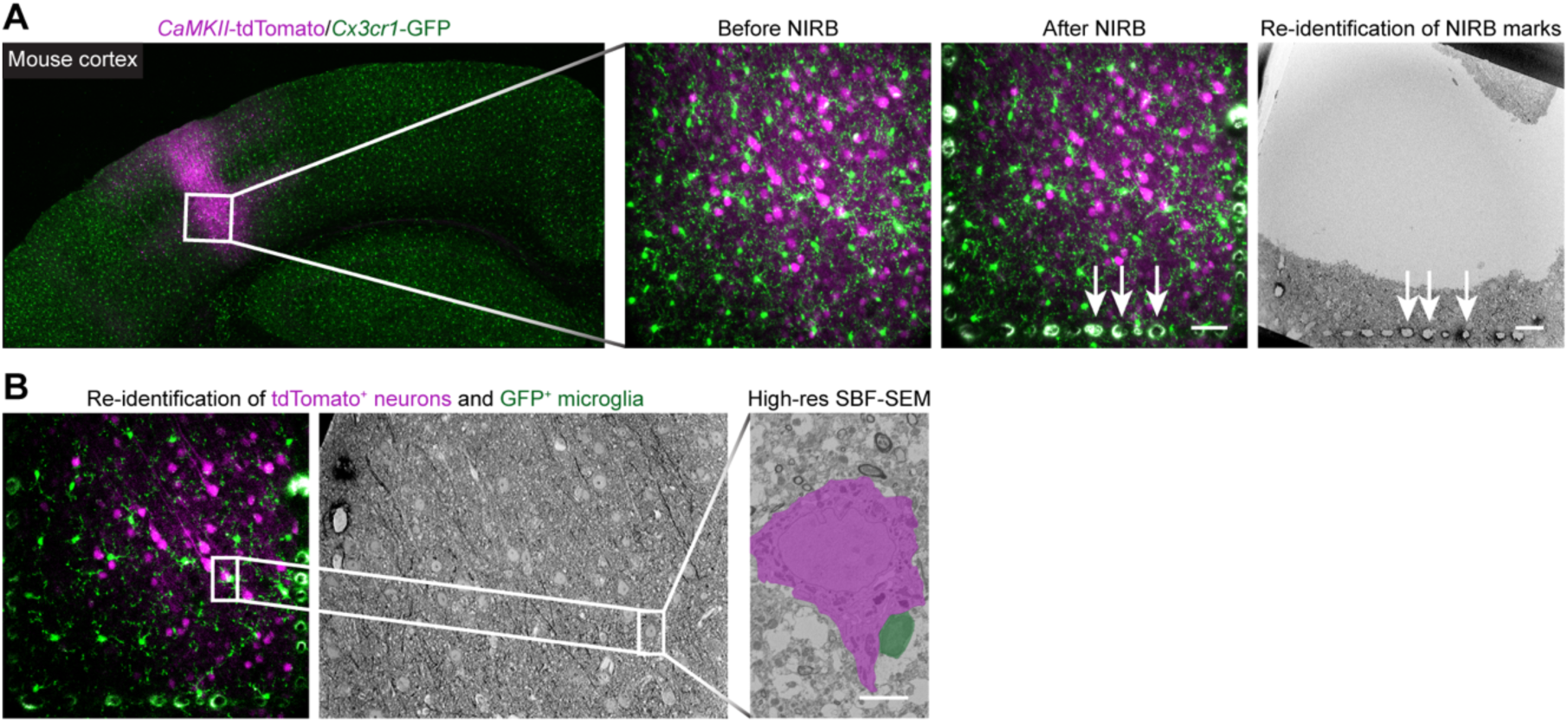
Workflow for examining membrane-membrane contact between satellite microglia and its associated neuron. (**A** and **B**) Fluorescence and EM images illustrating the stepwise identification of satellite microglia under SBF-SEM. *Cx3cr1^GFP/+^* mice received intracranial *AAV-CaMKII-tdTomato* injection to label cortical excitatory neurons (microglia, green; excitatory neurons, magenta). Brain sections were marked by near-infrared branding (NIRB) for relocation. NIRB marks were visible and matched under SBF-SEM (A). By combining NIRB marks with the distribution of nearby neuronal somata, the same satellite microglia and its contacting neuron were identified. Membrane-membrane contact was then examined by high-resolution SBF-SEM imaging (B). Scale bar: 5 μm (B) and 50 μm (A).

**Fig. S2.**
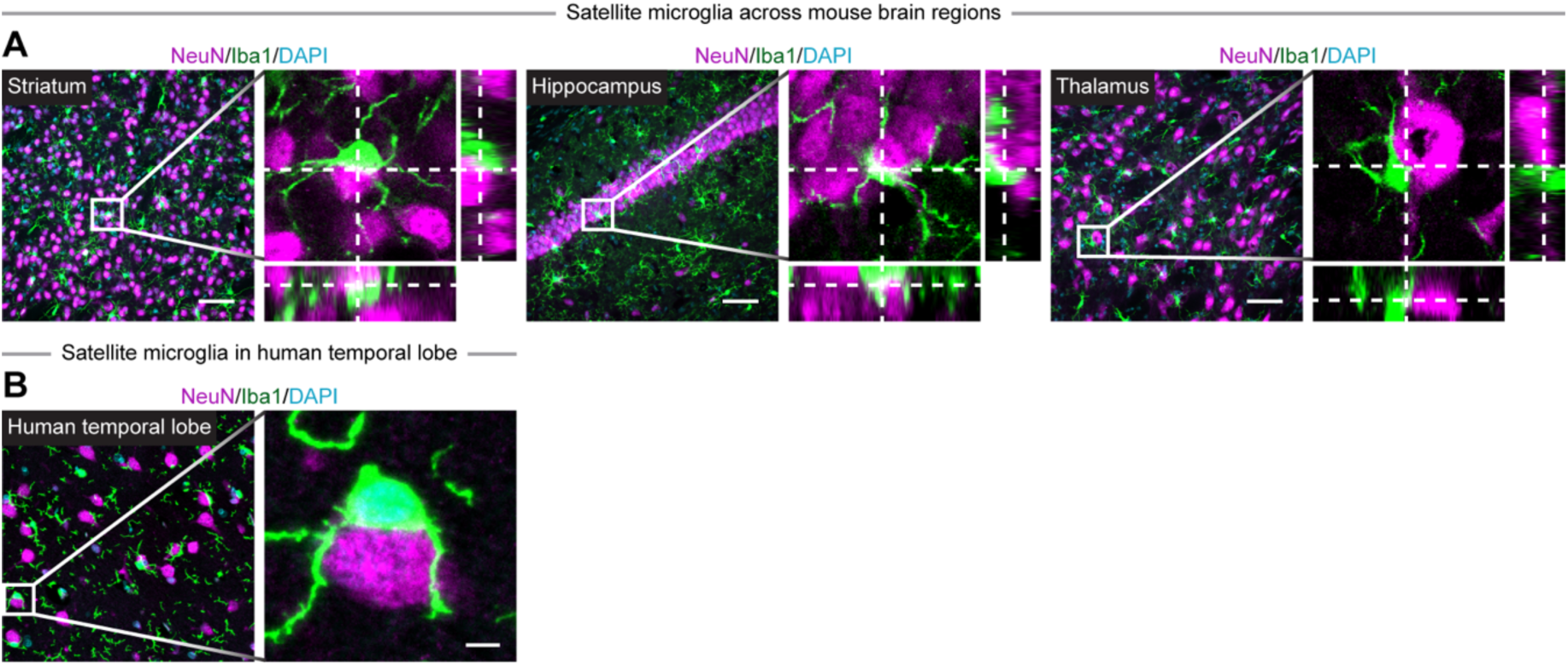
Satellite microglia-neuron soma interaction in mouse and human brains. (**A**) Immunostaining of satellite microglia (Iba1^+^, green) associated with neuronal somata (NeuN^+^, magenta) in the striatum, hippocampus, and thalamus. (**B**) Immunostaining (microglia, Iba1^+^, green; neurons, NeuN^+^, magenta) showing satellite microglia in the human temporal lobe. Scale bar: 5 μm (B) and 50 μm (A).

**Fig. S3.**
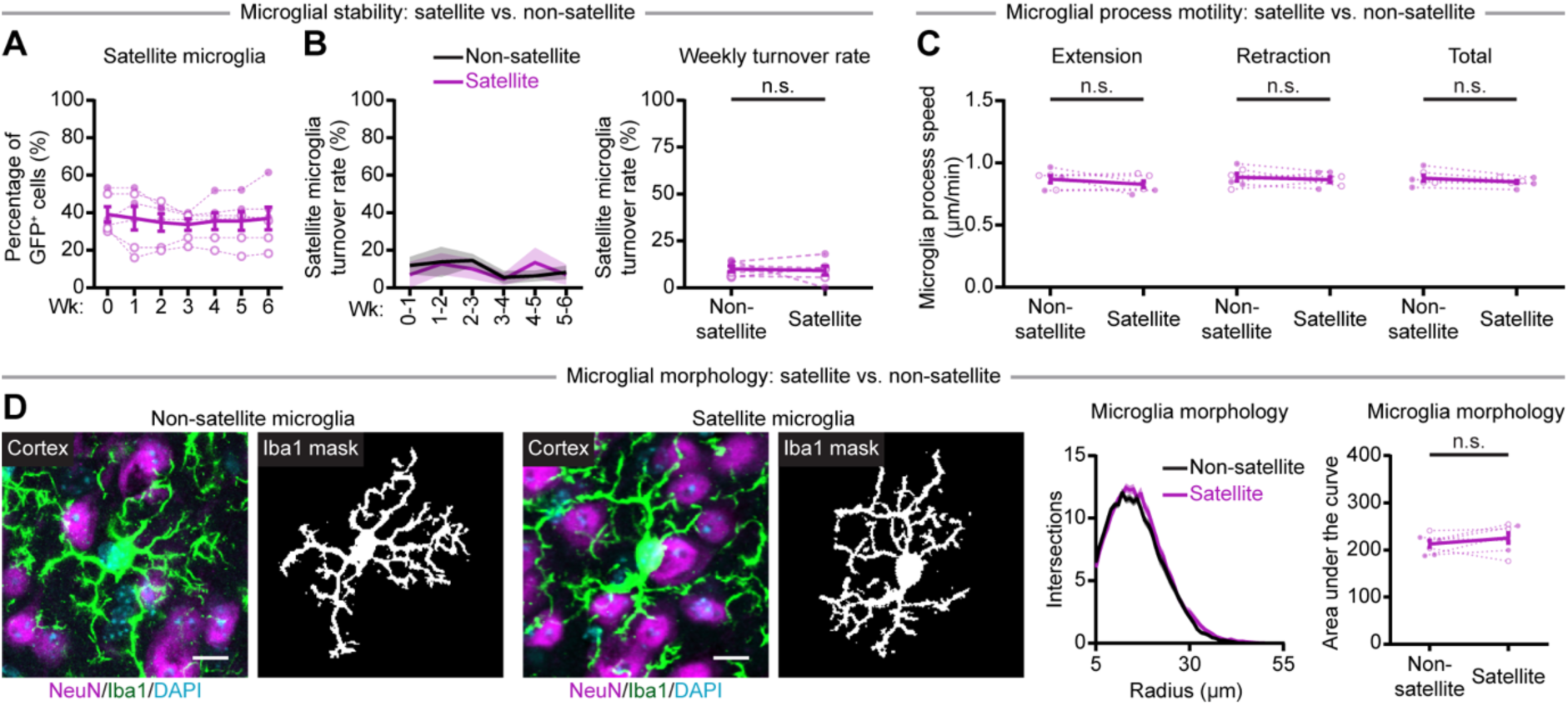
Satellite microglia and non-satellite microglia have similar stability and process dynamics. (**A** and **B**) Quantifications of long-term two-photon imaging revealing that satellite microglia consistently constituted approximately 40% of microglia (A), with a turnover rate of 9.11 ± 2.33% (B, *N* = 6). Satellite microglia and non-satellite microglia have similar weekly turnover rate. (**C**) Quantification of two-photon imaging of microglia demonstrating comparable process dynamics between satellite and non-satellite microglia (*N* = 6). (**D**) Immunostaining of non-satellite microglia (Iba1^+^, green), satellite microglia (Iba1^+^, green) and neurons (NeuN^+^, magenta) in the cortex. Quantifications of microglial morphology showing no significant differences between satellite and non-satellite microglia (*N* = 6). Each point indicates an individual mouse (solid dots: male; hollow dots: female). Data are presented as mean ± SEM. Paired *t*-test (B, C, and D). n.s.: not significant. Scale bar: 10 μm (D).

**Fig. S4.**
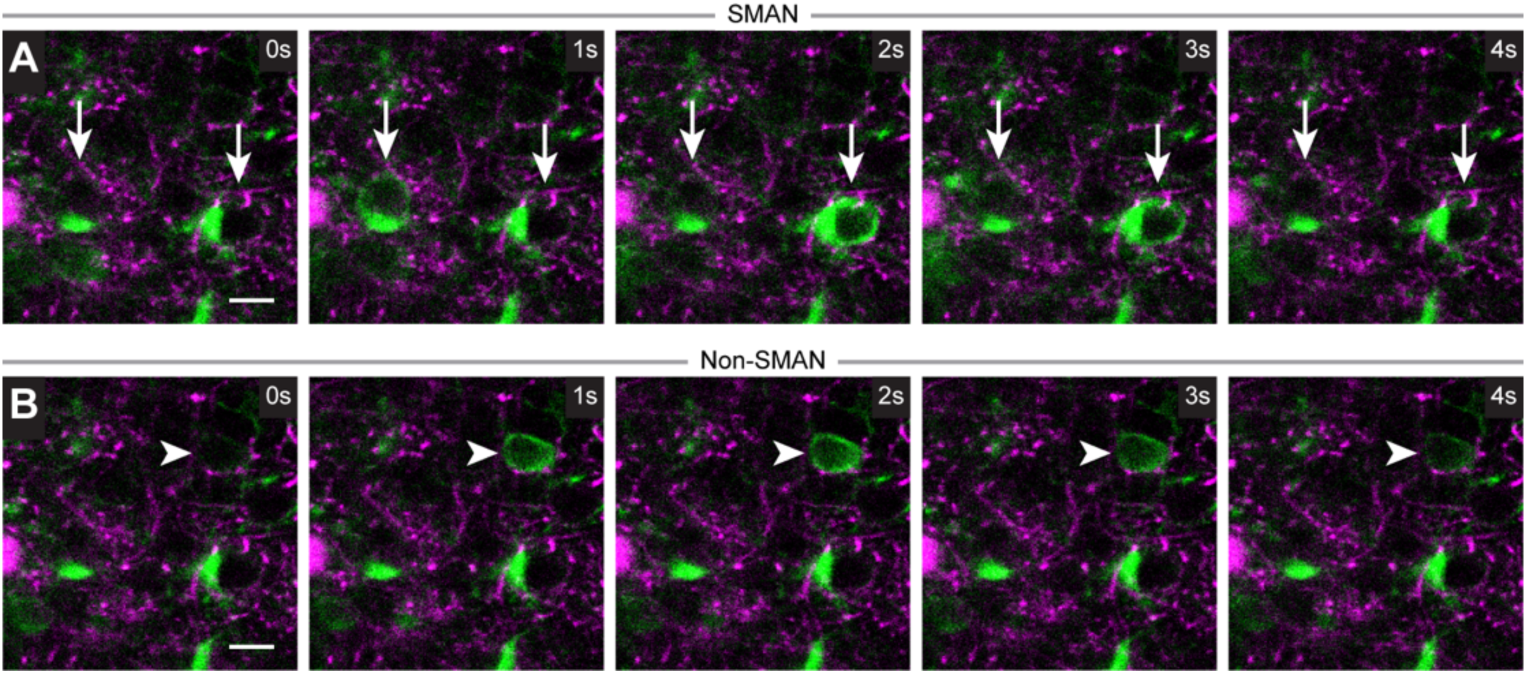
Two-photon imaging strategy to visualize inhibitory synaptic plasticity mediated by satellite microglia. (**A** and **B**) Two-photon images illustrating the identification of SMANs by the overlap between constant microglial GFP fluorescence and transient neuronal GCaMP6s signals (A, arrows), whereas non-SMANs showed only transient neuronal GCaMP6s signals (B, arrowheads). Scale bar: 10 μm.

**Fig. S5.**
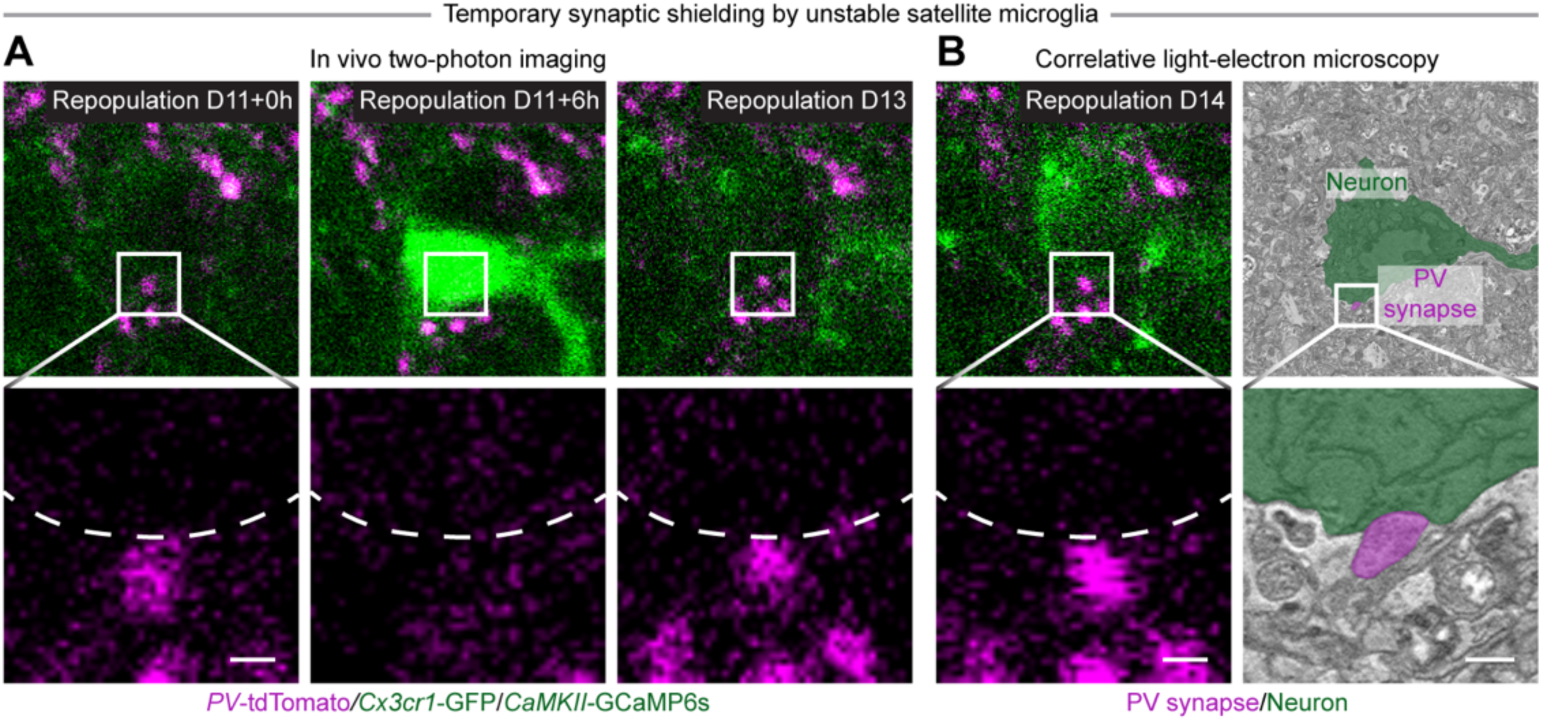
Unstable satellite microglia transiently displace PV^+^ puncta during repopulation. (**A**) Two-photon images illustrating transient synaptic shielding by unstable satellite microglia during repopulation. Repopulating microglia (green) transiently contacted the neuronal soma, during which the perineuronal synapse (magenta) was temporarily displaced (D11). Once the satellite microglia migrated away (D13), the synapse reestablished the contact. Dashed lines indicate neuronal somata. (**B**) Two photon and SBF-SEM images showing the same neuron (green) and inhibitory synapse (magenta). High resolution SBF-SEM imaging showing enriched synaptic vesicles were observed in transiently displaced inhibitory synapses (magenta). Scale bar: 1 μm (A and B).

**Fig. S6.**
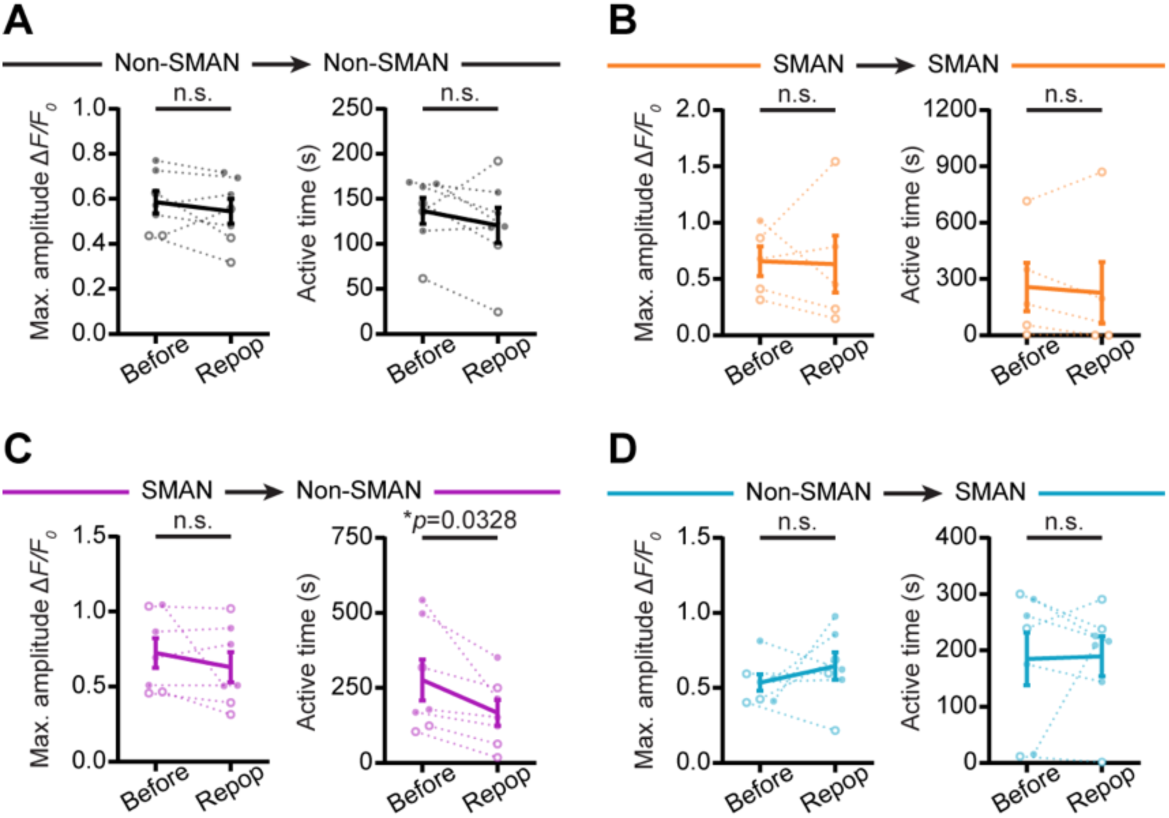
Effects of microglia redistribution on neuronal activity. (**A**) Comparison of Ca^2+^ activity in non-SMANs (non-SMAN → non-SMAN; *N* = 7, paired *t*-test) before and after microglia redistribution. Non-SMANs maintained similar maximum amplitude and active time. (**B**) Comparison of Ca^2+^ activity in SMANs (SMAN → SMAN; *N* = 5, not detected in 2 mice) before and after microglia redistribution. SMANs maintained similar maximum amplitude (Paired *t*-test) and active time (Wilcoxon test). (**C**) Quantification of Ca^2+^ activity in former SMANs (SMAN → non-SMAN; *N* = 7, paired *t*-test) showing no significant change in maximum amplitude but decreased active time. (**D**) Quantification of Ca^2+^ activity in newly formed SMANs (non-SMAN → SMAN; *N* = 7, paired *t*-test) showing no significant change in maximum amplitude and active time. Each point indicates an individual mouse (solid dots: male; hollow dots: female). Data are presented as mean ± SEM. n.s.: not significant.

**Fig. S7.**
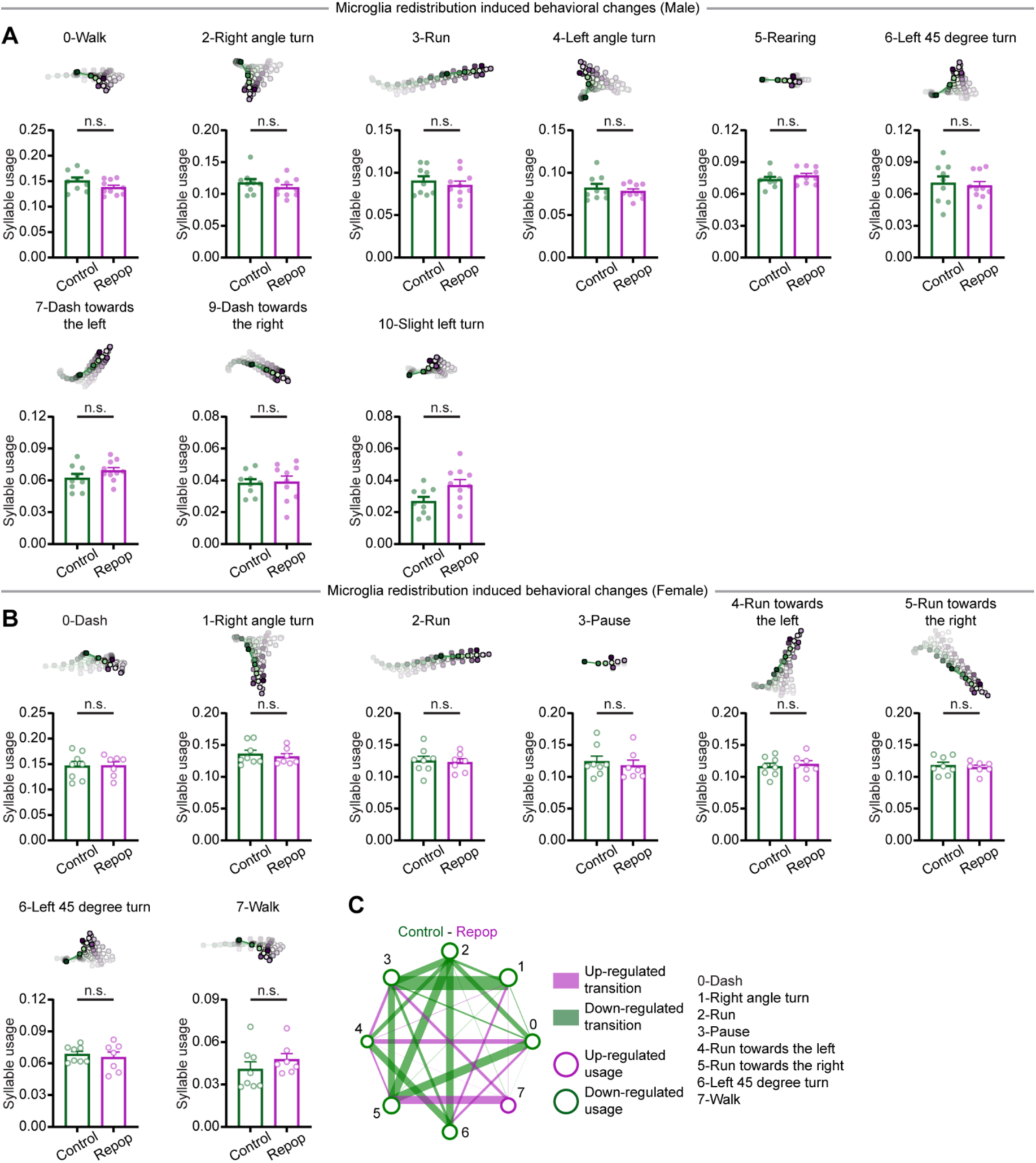
Additional comparisons of identified mouse behavior motifs. (**A**) Pose trajectories and quantifications of syllables in male mice. (**B**) Pose trajectories and quantifications of syllables in female mice. (**C**) Graph illustrating transition probability differences across syllables in female mice. Edge thickness reflects the magnitude of difference. Only syllables with usage > 0.03 are included in the analysis. Multiple unpaired *t*-tests with FDR correction (*p* < 0.1; A and B). n.s.: not significant.

**Movie S1. Satellite microglia form stable soma-soma interaction with neurons.**

Weekly time lapse two-photon imaging illustrating that some satellite microglia (Cx3cr1-GFP^+^, green; arrow) remained stably associated with the same neuronal soma (CaMKII-tdTomato^+^, magenta) for over 18 weeks. Scale bar: 20 μm.

**Movie S2. Satellite microglia associated neurons (SMANs) show higher neuronal activity.**

Time lapse two-photon imaging illustrating calcium activity of a SMAN (CaMKII-jRGECO1a, magenta; arrow) over a 10-minute period. Microglia are labeled with Cx3cr1-GFP (green). Scale bar: 20 μm.

**Movie S3. Satellite microglia dissociation enables the formation of perisomatic inhibitory synapses.**

Time lapse two-photon imaging illustrating the formation of perisomatic PV^+^ synapses (magenta) following dissociation of satellite microglia (green) at different Z-levels. Satellite microglia dissociation occurred by day 1 (D1). Neuronal somata are labeled with transient GCaMP6s signals (green). Lower panels show magnified views of the boxed regions in the upper panels, highlighting the formation of PV^+^ synapses. Scale bar: 1 μm.

**Movie S4. Ultrastructure of a newly formed perisomatic inhibitory synapse following satellite microglia dissociation.**

A 12.5 μm-thick stack of SBF-SEM images acquired at 50 nm step size, illustrating a newly formed inhibitory synapse (magenta) on the neuronal soma (green). The synapse formed at a site previously occupied by a satellite microglia. Scale bar: 2 μm.

**Movie S5. Formation of satellite microglia displaces perisomatic inhibitory synapses.**

Hourly time lapse two-photon imaging illustrating the displacement of perisomatic PV^+^ synapses (magenta) during the formation of satellite microglia (green) at different Z-levels. Neuronal somata are labeled with transient GCaMP6s signals (green). Lower panels show magnified views of the boxed regions in the upper panels, highlighting the displacement of PV^+^ synapses. Dashed lines outline neuronal somata. Scale bar: 1 μm.

**Movie S6. Microglia ablation and repopulation redistribute satellite microglia.**

Time lapse two-photon imaging illustrating microglial distribution (Cx3cr1-GFP^+^, green) before ablation (Before), during early repopulation (Repop D0-D13) and after complete repopulation (Repop D14, D21, and D28). Neurons are labeled with CaMKII-tdTomato (magenta). Arrows indicate former SMAN. Arrowheads indicate newly formed SMAN. Scale bar: 20 μm.

**Movie S7. Microglia ablation enables the formation of perisomatic inhibitory synapses.**

Daily time lapse two-photon imaging illustrating the formation of perisomatic PV^+^ synapses (magenta) following microglia ablation (PLX D1-14) at different Z-levels. Microglia are labeled by Cx3cr1-GFP (green). Neuronal somata are labeled with transient GCaMP6s signals (green). Lower panels show magnified views of the boxed regions in the upper panels, highlighting the formation of PV^+^ synapses. Dashed lines outline neuronal somata. Scale bar: 1 μm.

**Movie S8. Ultrastructure of newly formed inhibitory synapses after microglia ablation.**

A 17.5 μm-thick stack of SBF-SEM images acquired at 50 nm step size, illustrating newly formed inhibitory synapses (magenta) on the neuronal soma (green) following microglia ablation. The synapses formed at a site previously occupied by a satellite microglia. Scale bar: 1 μm.

**Movie S9. Unstable satellite microglia reversibly displace perisomatic inhibitory synapses during early microglia repopulation.**

Time lapse two-photon imaging illustrating the transient displacement of perisomatic PV^+^ synapses (magenta) by unstable satellite microglia (green) at different Z-levels. Neuronal somata are labeled with transient GCaMP6s signals (green). Lower panels show magnified views of the boxed regions in the upper panels, highlighting the rapid changes of PV^+^ synapses. Dashed lines outline neuronal somata. Scale bar: 1 μm.

**Movie S10. Ultrastructure of transiently displaced inhibitory synapses during early microglia repopulation.**

A 15 μm-thick stack of SBF-SEM images acquired at 50 nm step size, illustrating transiently displaced inhibitory synapses (magenta) that still maintain membrane-membrane contact with the neuronal soma (green). Scale bar: 2 μm.

**Movie S11. Formation of satellite microglia displaces perisomatic inhibitory synapses during microglia repopulation.**

Hourly time lapse two-photon imaging illustrating the displacement of perisomatic PV^+^ synapses (magenta) after microglia repopulation (Repop D13 and D14) at different Z-levels. Microglia are labeled with Cx3cr1-GFP (green). Neuronal somata are labeled with transient GCaMP6s signals (green). Lower panels show magnified views of the boxed regions in the upper panels, highlighting the displacement of PV^+^ synapses. Dashed lines outline neuronal somata. Scale bar: 1 μm.

**Movie S12. Ultrastructure of newly formed satellite microglia and displaced inhibitory synapse during microglia repopulation.**

A 15 μm-thick stack of SBF-SEM images acquired at 50 nm step size, illustrating a newly formed satellite microglia (orange) interposed between a neuronal soma (cyan) and an inhibitory synapse (magenta). The ultrastructure analysis highlights the physical displacement during the formation of satellite microglia. Scale bar: 1 μm.

**Movie S13. Redistribution of satellite microglia alters neuronal activity.**

Time lapse two-photon imaging illustrating calcium activity alterations of a former SMAN (CaMKII-jRGECO1a, magenta; arrow) and a newly formed SMAN (arrowhead). 10-minute movies were captured before microglia ablation (Before) and after complete repopulation (Repop). Microglia are labeled with Cx3cr1-GFP (green). Scale bar: 20 μm.

## References and Notes

1. A. Citri, R. C. Malenka, Synaptic plasticity: multiple forms, functions, and mechanisms. Neuropsychopharmacology 33, 18–41 (2008).

2. R. C. Paolicelli et al., Microglia states and nomenclature: A field at its crossroads. Neuron 110, 3458–3483 (2022).

3. S. Zhao, A. D. Umpierre, L. J. Wu, Tuning neural circuits and behaviors by microglia in the adult brain. Trends Neurosci 47, 181–194 (2024).

4. V. Duran Laforet, D. P. Schafer, Microglia: Activity-dependent regulators of neural circuits. Ann N Y Acad Sci 1533, 38–50 (2024).

5. P. del Río-Hortega, El “tercer elemento” de los centros nerviosos. I. La microglía en estado normal. Boletín de la Sociedad Española de Biología 8, 67–82 (1919).

6. O. Bakina, H. Kettenmann, C. Nolte, Microglia form satellites with diberent neuronal subtypes in the adult murine central nervous system. J Neurosci Res 100, 1105–1122 (2022).

7. V. Stratoulias, J. L. Venero, M. E. Tremblay, B. Joseph, Microglial subtypes: diversity within the microglial community. EMBO J 38, e101997 (2019).

8. K. Krukowski et al., Novel microglia-mediated mechanisms underlying synaptic loss and cognitive impairment after traumatic brain injury. Brain Behav Immun 98, 122–135 (2021).

9. Y. Wan et al., Microglial Displacement of GABAergic Synapses Is a Protective Event during Complex Febrile Seizures. Cell Rep 33, 108346 (2020).

10. T. Araki et al., Microglia induce auditory dysfunction after status epilepticus in mice. Glia 72, 274–288 (2024).

11. Z. Chen et al., Microglial displacement of inhibitory synapses provides neuroprotection in the adult brain. Nat Commun 5, 4486 (2014).

12. G. C. Brown, J. J. Neher, Microglial phagocytosis of live neurons. Nat Rev Neurosci 15, 209–216 (2014).

13. U. B. Eyo et al., P2Y12R-Dependent Translocation Mechanisms Gate the Changing Microglial Landscape. Cell Rep 23, 959–966 (2018).

14. A. Nimmerjahn, F. Kirchhob, F. Helmchen, Resting microglial cells are highly dynamic surveillants of brain parenchyma in vivo. Science 308, 1314–1318 (2005).

15. C. Cserep et al., Microglia monitor and protect neuronal function through specialized somatic purinergic junctions. Science 367, 528–537 (2020).

16. A. Badimon et al., Negative feedback control of neuronal activity by microglia. Nature 586, 417–423 (2020).

17. K. Haruwaka et al., Microglia enhance post-anesthesia neuronal activity by shielding inhibitory synapses. Nat Neurosci 27, 449–461 (2024).

18. T. F. Freund, I. Katona, Perisomatic inhibition. Neuron 56, 33–42 (2007).

19. A. Shapson-Coe et al., A petavoxel fragment of human cerebral cortex reconstructed at nanoscale resolution. Science 384, eadk4858 (2024).

20. C. Weinreb et al., Keypoint-MoSeq: parsing behavior by linking point tracking to pose dynamics. Nat Methods 21, 1329–1339 (2024).

21. N. Gutierrez-Castellanos, B. F. A. Husain, I. C. Dias, S. Q. Lima, Neural and behavioral plasticity across the female reproductive cycle. Trends Endocrinol Metab 33, 769–785 (2022).

22. S. Zhao et al., Chemogenetic activation of microglial Gi signaling decreases microglial surveillance and impairs neuronal synchronization. Sci Adv 11, eado7829 (2025).

23. A. Miyamoto et al., Microglia contact induces synapse formation in developing somatosensory cortex. Nat Commun 7, 12540 (2016).

24. D. P. Schafer et al., Microglia sculpt postnatal neural circuits in an activity and complement-dependent manner. Neuron 74, 691–705 (2012).

25. E. Favuzzi et al., GABA-receptive microglia selectively sculpt developing inhibitory circuits. Cell 184, 4048–4063 e4032 (2021).

26. A. Hashimoto et al., Microglia enable cross-modal plasticity by removing inhibitory synapses. Cell Rep 42, 112383 (2023).

27. Z. P. Chen et al., GABA-dependent microglial elimination of inhibitory synapses underlies neuronal hyperexcitability in epilepsy. Nat Neurosci 28, 1404–1417 (2025).

28. Y. Zou et al., Microglial pruning of glycinergic synapses disinhibits spinal PKCgamma interneurons to drive pain hypersensitivity in mice. Sci Transl Med 17, eadk8096 (2025).

29. P. T. Nguyen et al., Microglial Remodeling of the Extracellular Matrix Promotes Synapse Plasticity. Cell 182, 388–403 e315 (2020).

30. C. N. Parkhurst et al., Microglia promote learning-dependent synapse formation through brain-derived neurotrophic factor. Cell 155, 1596–1609 (2013).

31. L. J. Zhou et al., Microglia Are Indispensable for Synaptic Plasticity in the Spinal Dorsal Horn and Chronic Pain. Cell Rep 27, 3844–3859 e3846 (2019).

32. M. B. Dalva, A. C. McClelland, M. S. Kayser, Cell adhesion molecules: signalling functions at the synapse. Nat Rev Neurosci 8, 206–220 (2007).

33. K. Blinzinger, G. Kreutzberg, Displacement of synaptic terminals from regenerating motoneurons by microglial cells. Z Zellforsch Mikrosk Anat 85, 145–157 (1968).

34. S. Hebt, P. Jonas, Asynchronous GABA release generates long-lasting inhibition at a hippocampal interneuron-principal neuron synapse. Nat Neurosci 8, 1319–1328 (2005).

35. M. Capogna, P. E. Castillo, A. Mabei, The ins and outs of inhibitory synaptic plasticity: Neuron types, molecular mechanisms and functional roles. Eur J Neurosci 54, 6882–6901 (2021).

36. Y. K. Wu, C. Miehl, J. Gjorgjieva, Regulation of circuit organization and function through inhibitory synaptic plasticity. Trends Neurosci 45, 884–898 (2022).

37. H. C. Barron, T. P. Vogels, T. E. Behrens, M. Ramaswami, Inhibitory engrams in perception and memory. Proc Natl Acad Sci U S A 114, 6666–6674 (2017).

38. O. Marin, Interneuron dysfunction in psychiatric disorders. Nat Rev Neurosci 13, 107–120 (2012).

39. L. Verret et al., Inhibitory interneuron deficit links altered network activity and cognitive dysfunction in Alzheimer model. Cell 149, 708–721 (2012).

40. H. Wake, A. J. Moorhouse, S. Jinno, S. Kohsaka, J. Nabekura, Resting microglia directly monitor the functional state of synapses in vivo and determine the fate of ischemic terminals. J Neurosci 29, 3974–3980 (2009).

41. U. B. Eyo et al., Neuronal hyperactivity recruits microglial processes via neuronal NMDA receptors and microglial P2Y12 receptors after status epilepticus. J Neurosci 34, 10528–10540 (2014).

42. Y. U. Liu et al., Neuronal network activity controls microglial process surveillance in awake mice via norepinephrine signaling. Nat Neurosci 22, 1771–1781 (2019).

43. R. D. Stowell et al., Noradrenergic signaling in the wakeful state inhibits microglial surveillance and synaptic plasticity in the mouse visual cortex. Nat Neurosci 22, 1782–1792 (2019).

44. Y. You et al., Cell-specific IL-1R1 regulates the regional heterogeneity of microglial displacement of GABAergic synapses and motor learning ability. Cell Mol Life Sci 81, 116 (2024).

45. Y. Chen et al., Spatiotemporally selective astrocytic ATP dynamics encode injury information sensed by microglia following brain injury in mice. Nat Neurosci 27, 1522–1533 (2024).

46. N. J. Rothwell, G. N. Luheshi, Interleukin 1 in the brain: biology, pathology and therapeutic target. Trends Neurosci 23, 618–625 (2000).

47. D. Bishop et al., Near-infrared branding ebiciently correlates light and electron microscopy. Nat Methods 8, 568–570 (2011).

48. J. C. Fiala, Reconstruct: a free editor for serial section microscopy. J Microsc 218, 52–61 (2005).

49. J. Buchanan et al., Oligodendrocyte precursor cells ingest axons in the mouse neocortex. Proc Natl Acad Sci U S A 119, e2202580119 (2022).

50. A. Mathis et al., DeepLabCut: markerless pose estimation of user-defined body parts with deep learning. Nat Neurosci 21, 1281–1289 (2018).

